# Mitophagy promotes metabolic reprogramming to enhance keratinocyte migration via ANGPTL4 during wound healing

**DOI:** 10.64898/2025.12.03.692043

**Authors:** Matthew Hunt, Nuoqi Wang, Monica Torres, Jenna Villman, Ilkka Paatero, Shannon Hinch, Gustavo Urbano-Quispe, Margarita Chatzopoulou, Etty Bachar-Wikström, Jakob D Wikström

## Abstract

Mitochondrial function and quality control is emerging as a key regulator of keratinocyte migration in both wounding and non-wound healing contexts, yet the cellular mechanisms that support this process are incompletely understood. In this study, using single-cell RNA sequencing data of human wounded tissue we identified a distinct population of migrating keratinocytes marked by the high expression of the mitophagy regulator *BNIP*3 during the proliferative stage of wound healing. Pharmacological induction of mitophagy with Urolithin A accelerated keratinocyte migration *in vitro* as well as keratinocyte function and regeneration in aged zebrafish, whilst RNA sequencing of primary human keratinocytes revealed the transcriptional upregulation of *ANGPTL4* in Urolithin-A treated cells. Mechanistically, Urolithin A increased metabolic switching to a more glycolytic phenotype, leading to AKTGSK3 pathway activity and FOSL1-mediated *ANGPTL4* transcription, ultimately promoting keratinocyte migration through enhanced laminin-332 production. Overall, our findings uncover a novel role for mitophagy in promoting keratinocyte migration during wound repair, and demonstrate that pharmacologically enhancing mitophagy promotes regenerative epithelial responses by enhancing FOSL1-mediated ANGPTL4 signalling through the modulation of metabolic switching. These insights significantly expand the understanding of the role of mitophagy on keratinocyte function during wound healing, linking mitophagy to metabolic adaption in keratinocytes, and provide a mechanistic basis for targeting mitophagy or downstream genes in wound healing therapies.

## Introduction

In response to cutaneous injury, the conserved multi-step process of wound healing is required to restore skin barrier structure and function^1^. Would healing consists of the four concurrent and overlapping phases of: haemostasis, inflammation, proliferation, and tissue remodelling^2^. In the event of dysregulation in this wound healing process, chronic wounds (CW) form. CWs are associated with a significant decrease in both morbidity and quality of life^3^ and account for approximately 2-4% of healthcare budgets worldwide^4^. At present, there are few validated treatments for CWs, owed in part to the multifaceted nature and large variation in pathophysiology between patients, as well as the fact that significant gaps remain regarding the underlying cellular and molecular mechanisms of wound healing and CW formation^5^.

Keratinocytes play vital roles in the wound healing process, in particular with regards to reepithelialisation, where wound edge keratinocytes actively proliferate and migrate in order to establish the epithelial barrier and thus aid in healing^6^. In addition, keratinocytes produce numerous factors which promote angiogenesis as well as extracellular matrix (ECM) remodelling and formation^6, 7^.

Mitochondria facilitate important roles in the various cell types involved in dermal wound healing, including modulating reactive oxygen species (ROS) levels and signalling^8^, and regulating metabolic reprogramming in order to facilitate macrophage activation^9^, as well as in both fibroblast and keratinocyte migration^10^. In addition, mitochondrial function is required in order for angiogenesis^11^ and ECM remodelling^12^ during wound healing. Mitochondrial autophagy (mitophagy) is the selective sequestration and clearance of mitochondria by autophagy, and effective regulation of mitophagy is essential for both the functional integrity of the mitochondrial network and cell homeostasis^13, 14^. There are numerous described pathways of mitophagy, which is triggered in response to stressors such as hypoxia, cellular starvation, or mitochondrial damage^15^.

Whilst it has been demonstrated that mitophagy promoted keratinocyte migration and proliferation during wound healing through the degradation of p-MAP4^16^, and that in non-wound healing contexts BNIP3-induced mitophagy accelerated keratinocyte migration^17, 18^, and separately that both BNIP3- and NIX-mediated mitophagy play important roles in keratinocyte differentiation in the epidermis^19, 20, 21^, at present, the specific cellular and molecular roles of mitophagy in wound healing, and in particular keratinocyte function during wound healing, are incompletely understood^22^. In this study, through the re-analysis of human wound healing single-cell RNA-sequencing (scRNAseq) data we show that ubindependent mitophagy, particularly BNIP3-mediated mitophagy, is upregulated in migrating basal and spinous keratinocytes during the proliferation stage of wound healing, and that mitophagy is upregulated and promotes migration in keratinocytes *in vitro*.

Pharmacologically inducing mitophagy through Urolithin A (UroA) treatment enhanced the transcription of *ANGPTL4*, mediated through increases in glycolysis and decreases in mitochondrial ROS levels, leading to AKT-GSK3 pathway activation and FOSL1 transcription factor activity. Increased ANGPTL4 ultimately accelerated keratinocyte migration through upregulating laminin-332 production.

## Results

### Ub-independent mitophagy is upregulated in keratinocytes at the proliferation stage of human wound healing

To investigate the expression of mitophagy markers in keratinocytes during normal wound healing, we analysed single-cell RNA sequencing (scRNAseq) data of human wound healing which included day 0 (intact), day 1, day, and day 30 post wounding samples, representing the different stages of wound healing respectively. Following clustering, which determined there to be 9 subclusters of keratinocytes (**Figure 1A**), we investigated the differential expression (DE) of both ub-dependent and ub-independent mitophagy markers at days 1, 7, and 30 compared to day 0 in basal migration and spinous migration subclusters. Here, the expression of numerous ub-dependent mitophagy markers such as *NIX* and *FUNDC1*, as well as ub-dependent markers such as *PRKN* and *OPTN* were downregulated at day 1 compared to day 0, whilst at day 7, several ub-independent markers were upregulated, particularly *BNIP3* (**Figure 1B**). The DE of both ub-dependent and ub-independent markers was less pronounced in day 30 vs day 0. Next, we then examined the expression of *BNIP3* in all 9 keratinocyte subclusters at the four time points. Here, whilst the *BNIP3* expression was high in the granular subcluster at day 0 and subsequently declined, *BNIP3* expression increased to the greatest extent in both basal migration and spinous migration subclusters at day 7 (**Figure 1C**). This was validated through analysis of BNIP3 expression in basal and spinous migration from spatial transcriptomics data of human wound healing (**Supplementary Figure 1C**), which demonstrated a significantly greater number of BNIP3expressing spots for these cell types (**Supplementary Figure 1D**). Altogether, these results suggested an important role for BNIP3 mitophagy in keratinocyte migration during the proliferation stage of wound healing.

**Figure 1.**
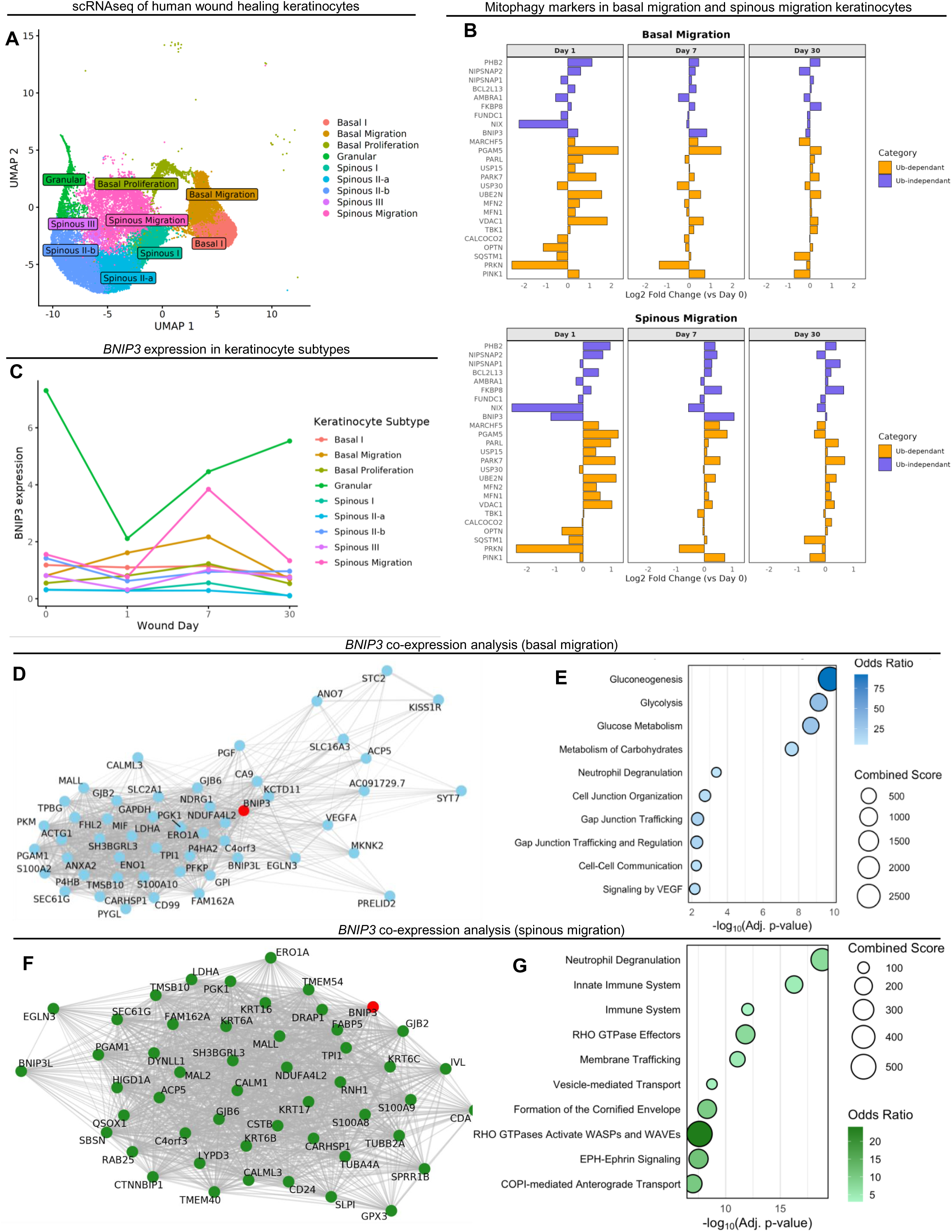
*BNIP3* is upregulated in migrating keratinocytes during human wound healing. (**A**) UMAP of keratinocyte clusters from human acute wound healing samples (**B**) Graphs showing Log_2_ fold change in the expression of ub-dependant (orange) and ub-independent (blue) genes in day 1, 7, and 30 vs intact within basal and spinous migration clusters. (**C**) Expression of *BNIP3* at wound days in the 9 keratinocyte clusters. (**D**) Map of top 50 genes co-regulated with *BNIP3* (red) in basal migration keratinocytes at day 7. (**E**) Pathway enrichment analysis of top co-regulated genes with *BNIP3* in basal migration keratinocytes at day 7. (**F**) Map of top 50 genes co-regulated with *BNIP3* (red) in spinous migration keratinocytes at day 7. (**G**) Pathway enrichment analysis of top co-regulated genes with *BNIP3* in spinous migration keratinocytes at day 7.

Next, to investigate the functional role of BNIP3 in basal and spinous migration keratinocytes at day 7, we then performed gene co-expression and pathway enrichment analysis of the top co-expressed genes. In basal migration keratinocytes, gene co-expression (**Figure 1D**) and enrichment analysis demonstrated a correlation of *BNIP3* with genes involved in glycolytic metabolism, as well as cell and gap junction organization (**Figure 1E**). In spinous migration keratinocytes, gene co-expression (**Figure 1F**) and enrichment analysis revealed a correlation of *BNIP3* with innate immune response regulation, as well as ‘RHO GTPase Effectors’, ‘Membrane Trafficking’, and ‘Formation of the Cornified Envelope’, among others (**Figure 1G**). This suggested a role of BNIP3 in glycolytic metabolism and junction regulation in basal migration keratinocytes, and that of immune response modulation and broad migration mechanisms in spinous migration keratinocytes, all collectively important for keratinocyte migration during the proliferation stage of wound healing.

### Mitophagy is upregulated in migrating keratinocytes *in vitro*

Following the demonstration that mitophagy marker gene expression was upregulated in keratinocytes at the proliferation stage of wound healing, particularly in migrating keratinocytes, we next investigated whether mitophagy was increased in migrating keratinocytes *in vitro*. Firstly, sequential daily live cell imaging of mitochondria and lysosomes in primary human epidermal keratinocytes (HEKa) seeded with a scratch wound (**Figure 2A-B**), demonstrated that the number of mitophagy events – as determined by the overlapping of mitochondria and lysosomal puncti – was increased in ‘wound edge’ and ‘migrating’ compared to ‘intact’ HEKa at 0, 24, and 48 hours respectively (**Figure 2C**). There was no significant difference at 72 and 96 hours post scratching, when the cells had fully migrated. In addition, mitochondrial mass was decreased in wound edge and migrating HEKa at 0, 24, and 48 hours when compared to stationary (**Figure 2D**), whilst mitochondria also had significantly decreased form factor and aspect ratio in wound edge and migrating HEKa at 24 hours (**Supplementary Figure 2A-B**). Overall, these results suggested that migrating HEKa had higher levels of mitophagy, decreased mitochondrial mass, and a more fragmented mitochondrial phenotype when compared to non-migrating HEKa.

**Figure 2.**
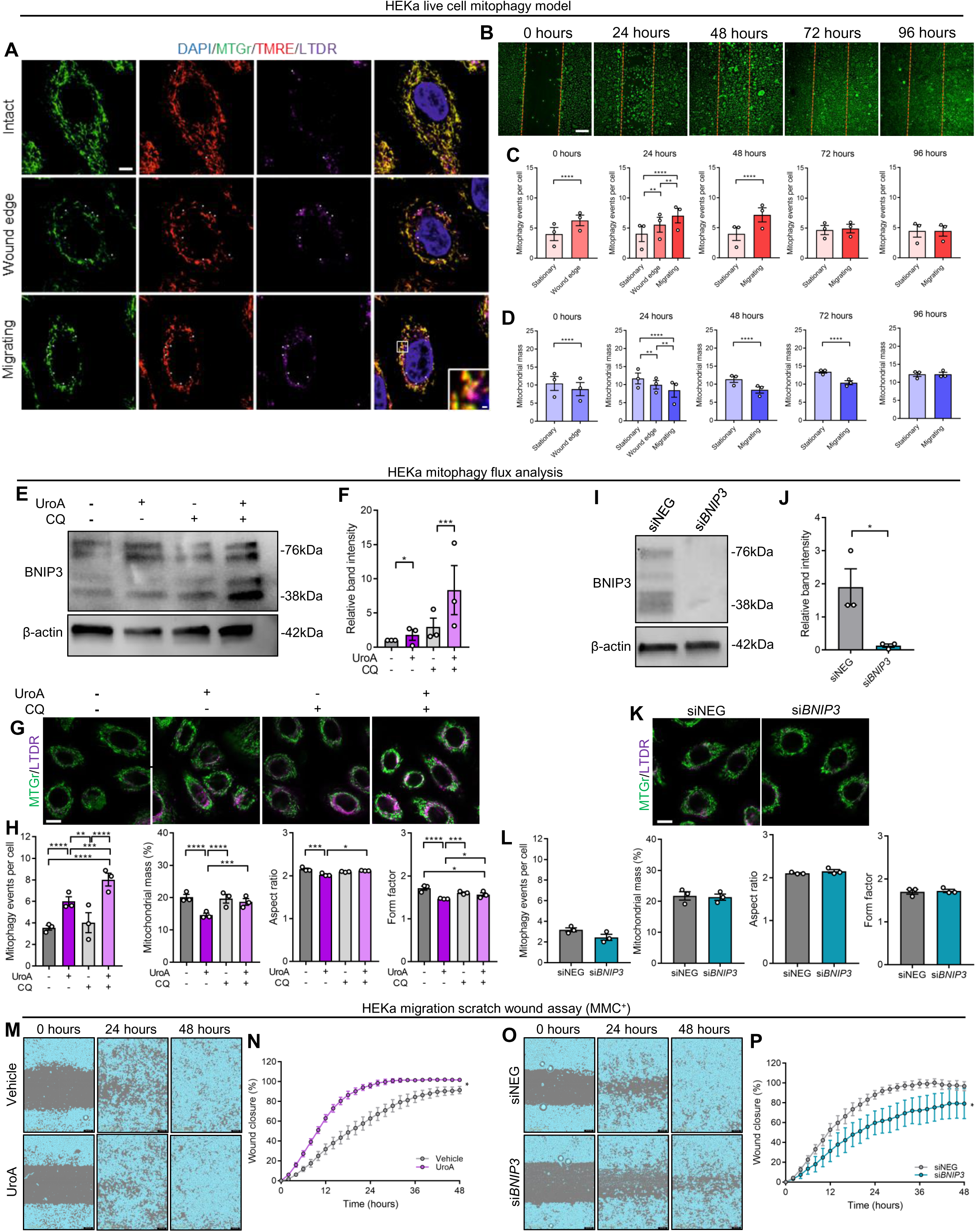
Mitophagy promotes migration in keratinocytes *in vitro*. (**A**) Representative fluorescence images of intact, wound edge, and migrating HEKa stained with DAPI, MTGr, TMRE, and LTDR. Arrows depict colocalised lysosomes and mitochondria, representing mitophagy events. Scale bars = 10µm (full size), 0.5µm (zoom). (**B**) Representative fluorescence images of HEKa at 0, 24, 48, 72, and 96 hours in the live cell wounding assay. Orange lines represent the wound edge. Scale bar = 200µm. I = intact; WE = wound edge; M = migratory. (**C-D**) Quantification (mean ± SEM) of (**C**) co-localised mitophagy events and (**D**) mitochondrial mass. Two-way ANOVA or students t-test, **** = p < 0.0001; *** = p < 0.001; ** = p < 0.01. N = 30 cells in 3 separate biological replicates within each condition. Each dot represents the mean of an individual biological replicate. (**E-F**) Representative immunoblot and (**F**) quantification (mean ± SEM) of BNIP3 in WT HEKa. Two-way ANOVA, *** = p < 0.001, * = p < 0.05. Each dot represents an individual biological replicate. (**I-J**) Quantification (mean ± SEM) of mitophagy events per cell, mitochondrial mass, aspect ratio, and form factor in WT HEKa. Two-way ANOVA, **** = p < 0.0001; *** = p < 0.001; ** = p < 0.01, * = p < 0.05. N = 15 cells in 3 separate biological replicates within each condition. Each dot represents the mean of an individual biological replicate. (**G**) Representative fluorescence images of MTGr and LTDR staining in WT HEKa. Scale bar = 100µm. (**I**) Representative immunoblot and (**J**) quantification (mean ± SEM) of BNIP3 in siNEG and si*BNIP3* HEKa. Students t-test, * = p < 0.05. Each dot represents an individual biological replicate. (K) Representative fluorescence images of MTGr and LTDR staining in siNEG and si*BNIP3* HEKa. Scale bar = 100µm. (L) Quantification (mean ± SEM) of mitophagy events per cell, mitochondrial mass, aspect ratio, and form factor in siNEG and si*BNIP3* HEKa. Students t-test, **** = p < 0.0001; *** = p < 0.001; ** = p < 0.01, * = p < 0.05. N = 15 cells in 3 separate biological replicates within each condition. Each dot represents the mean of an individual biological replicate. (**M-N**) Representative images and (**N**) quantification (mean ± SEM) of scratch assay migration following Mitomycin C treatment in vehicle and UroA-treated HEKa. Students ttest, * = p < 0.05. N = 3 biological repeats containing at least 2 technical repeats. (**O-P**) Representative images and (**P**) quantification (mean ± SEM) of scratch assay migration following Mitomycin C treatment in siNEG and si*BNIP3* HEKa. Students t-test, * = p < 0.05. N = 3 biological repeats containing at least 2 technical repeats.

We then sought to determine whether pharmacologically inducing mitophagy using UroA accelerated migration in HEKa. Firstly, mitophagy flux western blot analysis using the autophagy inhibitor chloroquine (CQ) demonstrated that UroA significantly increased BNIP3 levels (**Figure 2E-F**), and live cell imaging showed a significant increase in mitophagy events in UroA-treated HEKa (**Figure 2G-H**). As *BNIP3* levels were significantly increased in both scRNAseq analysis of migrating keratinocytes, as well as through UroA treatment, we next performed siRNA knockdown (KD) of *BNIP3* (si*BNIP3*) (**Figure 2I-J**), and confirmed that mitophagy events were significantly decreased in si*BNIP3* HEKa when compared to siNEG (**Figure 2K-L**). Finally, scratch wound migration analysis demonstrated firstly that UroA treatment significantly increased HEKa migration (**Figure 2M-N**), whilst HEKa migration was significantly decreased in si*BNIP3* HEKa compared to siNEG (**Figure 2O-P**), indicating a role of BNIP3-mediated mitophagy in accelerating keratinocyte migration.

### Urolithin A increases regeneration in aged zebrafish caudal fin model

In order to validate the beneficial effects of UroA *in vivo*, we next investigated caudal fin regeneration in aged (158-164 weeks) zebrafish treated with UroA or vehicle (**Figure 3A**). Here, quantification of caudal fin regeneration at days 4 and 8 after amputation (dpa) (**Figure 3B**) revealed a significant increase in caudal fin regeneration at both days in the UroA-treated zebrafish when compared to vehicle (**Figure 3C**). Determination of zebrafish body weight then revealed no significant difference in body weight between the two groups (**Figure 3D**), whilst toxicology and behavioural tests in zebrafish embryos treated with UroA (**Supplementary Figure 3**) demonstrated that UroA was not toxic to the zebrafish. Assessment of bony ray morphology demonstrated no difference between vehicle and UroA-treated zebrafish (**Figure 3E-G**). Finally, through RT-qPCR analysis of caudal fin biopsies, we then investigated the expression of markers known to be important in keratinocyte function during wound healing – *krt1, krt4,* and *krt8.* Here, expression of both *krt4* and *krt8* was significantly higher in UroA-treated zebrafish caudal fin sections when compared to vehicle at 4 dpa (**Figure 3H**). Overall, these results demonstrated the beneficial effect of UroA on keratinocyte function during regeneration processes. As such, we then focused our investigations in to exploring the underlying cellular and molecular mechanisms as to how mitophagy accelerates keratinocyte migration.

**Figure 3.**
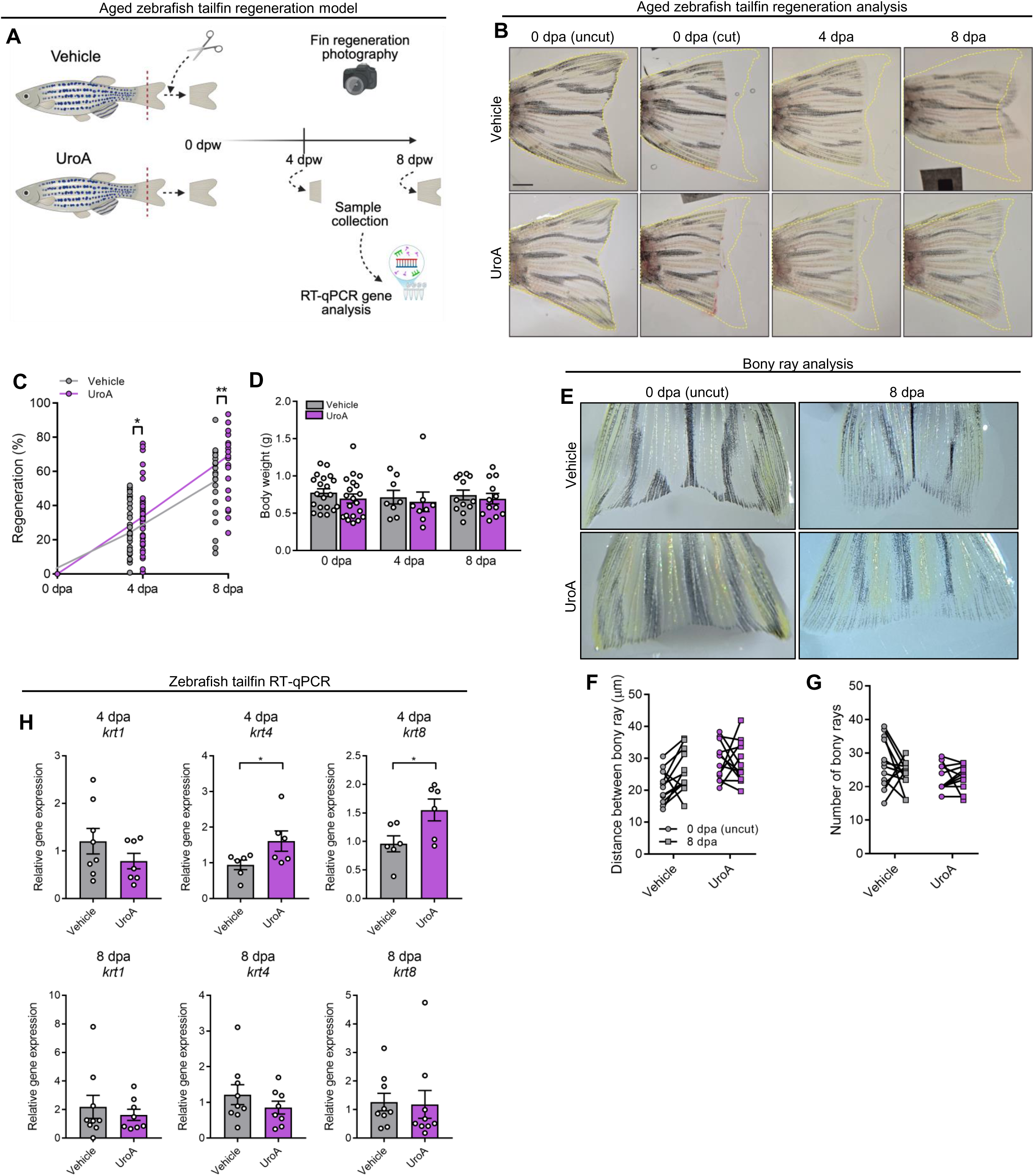
UroA promotes regeneration in aged zebrafish tailfin regeneration model. (**A**) Schematic representation of the experimental outline. (**B**) Representative images of zebrafish tailfins at uncut 0 days post amputation (dpa), amputated 0 dpa, 4 dpa, and 8 dpa. Orange line represents tailfin at 0 dpa (uncut). Scale bar = 1mm. (**C**) Quantification (mean ± SEM) of percentage tailfin regeneration. Two-way ANOVA, ** = p < 0.01, * = p < 0.05. Each dot represents an individual zebrafish at the respective time points (n = 24). (**D**) Quantification (mean ± SEM) of body weight (g). Two-way ANOVA. Each dot represents an individual zebrafish at the respective time points (n = 24). (**E**) Representative images of zebrafish tailfins and bony rays at 0 dpa (uncut) and 8 dpa. (**F-G**) Quantification (mean ± SEM) of (**F**) distance between bony rays (µm) and (**G**) number of bony ray branches at 0 dpa (uncut) (circle) and 8 dpa (square). Two-way ANOVA. Each dot represents an individual zebrafish (n = 12). (**H**) Quantification (mean ± SEM) of *krt1, krt4,* and *krt8* gene levels in vehicle and UroA treated zebrafish tailfins at 4 dpa and 8 dpa. Students t-test, * = p < 0.05. Each dot represents an individual zebrafish (n = 24).

### RNAseq analysis of UroA treatment in HEKa

To profile the effect of UroA treatment on gene expression, both vehicle- and UroA-treated HEKa underwent bulk RNAseq gene expression analysis (**Figure 4A**). Here, DE (**Figure 4B**) and enrichment analysis (**Figure 4C**) demonstrated a significant upregulation of genes involved in pathways associated with keratinocyte migration, including ‘Laminin Interactions’, ‘Non-integrin Membrane-ECM interactions’, ‘Collagen Formation’, ‘RHO GTPase Cycle’, and ‘Extracellular Matrix Organization’ in UroA-treated HEKa compared to vehicle. We then performed a series of gene KD experiments targeting the 20 most upregulated genes (**Figure 4D**) to identify those exerting the greatest impact on keratinocyte migration. Here, KD of several genes including *ANGPTL4, DERL3, CSF3, COL8A1, MRGPRX3, IL1RL1,* and *MYCBPAP* significantly decreased migration when compared to siNEG HEKa, whilst surprisingly, KD of *FAM196B* significantly increased migration (**Figure 4E**). RT-qPCR analysis confirmed the decrease in respective gene expression in HEKa (**Supplementary Figure 4A**), as well as an increase in *angptl4* at 8dpa in zebrafish caudal fins (**Figure 4G**). As such, we decided to focus subsequent studies on the genes in which KD decreased migration the to the greatest extent – *ANGPTL4, DERL3,* and *CSF3* (**Figure 4F**).

**Figure 4.**
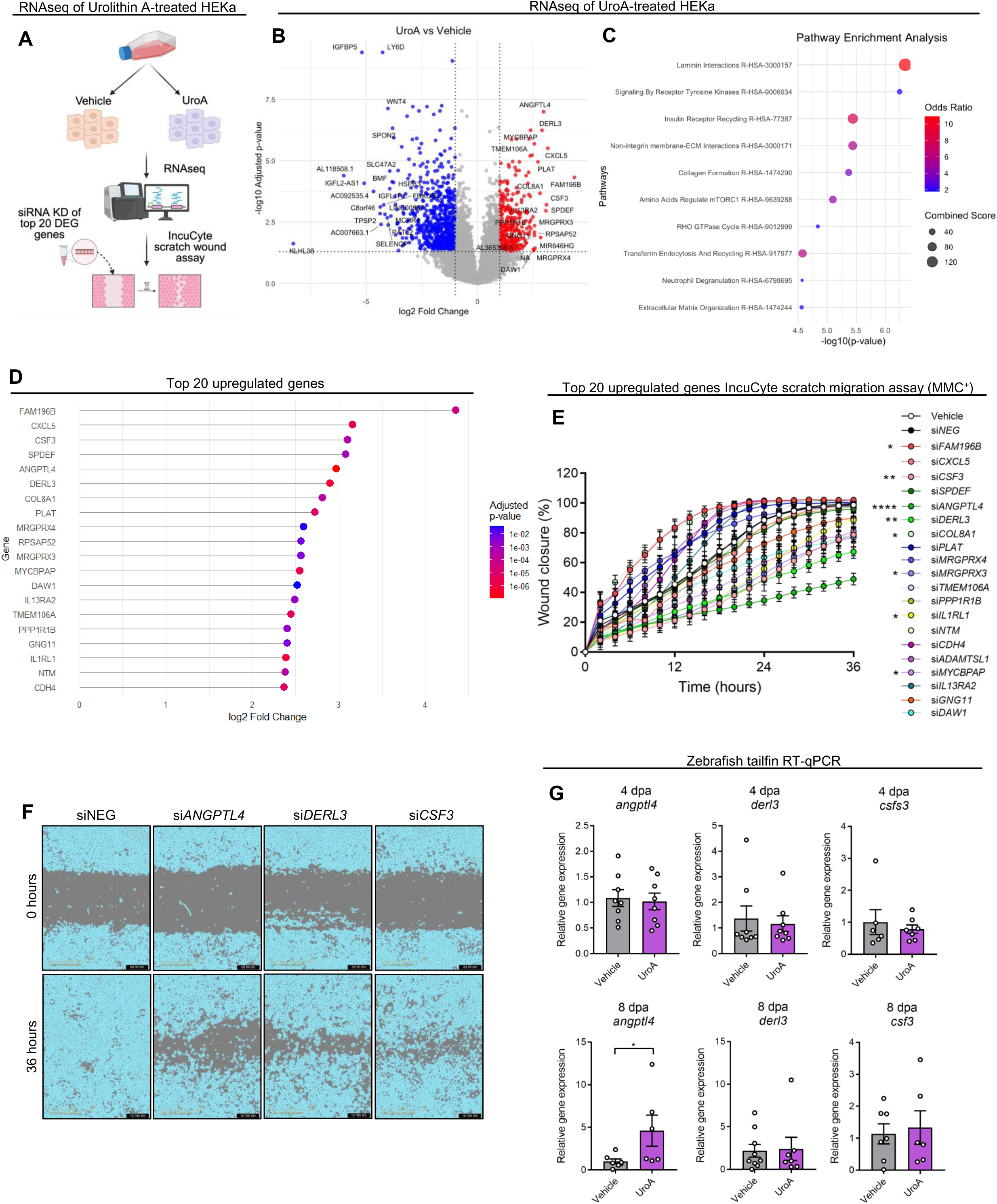
Gene expression analysis of UroA treatment. (**A**) Schematic representation of the experimental outline. (**B**) Volcano plot depicting DEG quantification of UroA vs vehicle HEKa gene expression. Genes significantly upregulated in UroA HEKa (red) and significantly downregulated (blue) compared to vehicle. (**C**) Gene enrichment analysis showing the 10 top significantly upregulated pathways in UroA treated HEKa from Reactome database. (**D**) Bubble plot depicting the log2 fold change and adjusted p-value of the top 20 significantly upregulated genes in UroA-treated HEKa. Higher adjusted p value is coloured red, lower adjusted p value is coloured bluer. (**E**) Quantification (mean ± SEM) of wound closure percentage. Two-way ANOVA, **** = p < 0.0001; ** = p < 0.01, * = p < 0.05. Each dot represents the mean of at least 2 technical repeats. (**F**) Representative images of siNEG, si*ANGPTL4,* si*DERL3*, si*CSF3* HEKa at 0 and 36 hours from scratch wound assay. (**G**) Quantification (mean ± SEM) of *angptl4, derl3,* and *csf3* gene expression in zebrafish tailfins 4 dpa and 8 dpa. Students t-test, * = p < 0.05. Each dot represents an individual zebrafish (n = 24).

### UroA promotes *ANGPTL4* expression to drive keratinocyte migration

We next sought to determine whether the UroA-induced increase in HEKa migration was mediated through *ANGPTL4, DERL3,* or *CSF3.* Here, KD of the three genes significantly decreased migration when compared to siNEG HEKa, but whilst treatment of si*DERL3* and si*CSF3* with UroA significantly increased their migration when compared to vehicle-treated siNEG, implying that UroA increased migration regardless of the KD of these two genes, indicating they do not contribute to UroA-action on keratinocytes migration. However, there was no significant difference between si*ANGPTL4* and si*ANGPTL4* + UroA cells (**Figure 5AD**) which suggested that the UroA-mediated increase in *ANGPTL4* drives HEKa migration, and the importance of *DERL3* and *CSF3* in keratinocyte migration could be explained by other mechanisms. In addition, UroA treatment significantly increased siNEG migration when compared to vehicle-treated siNEG (**Figure 5A-D**) and overexpression (OE) of *ANGPTL4* significantly increased migration when compared to pCMV control vector (**Supplementary Figure 5A-B**). Validating the role of *ANGPTL4* in keratinocyte migration during wound healing, immunofluorescence staining of ANGPTL4 in wounded tissue from aged mice demonstrated the significant increase in ANGPTL4 protein levels at the leading edge in UroA-treated mice compared to vehicle at 5 days post wounding (dpw) (**Figure 5E-F**), whilst assessment of *ANGPTL4* expression in both basal migration and spinous migration cells from the human scRNAseq data demonstrated the increase at day 7 (**Figure 5G**). Finally, we performed a rescue experiment in which transfection of both *siANGPTL4* followed by *ANGPTL4* OE and then treatment with UroA significantly increased HEKa migration when compared to both si*ANGPTL4* as well as *siANGPTL4* + UroA (**Figure 5H-I**), further suggesting that UroA-mediated migration acceleration is mediated through ANGPTL4. RTqPCR analysis confirmed the efficiency of both KD and OE, as well as double transfection, and although not significant, there was an almost double fold change inn *ANGPTL4* expression in double transfected + UroA when compared to vector control (**Supplementary Figures 5D-E**).

**Figure 5.**
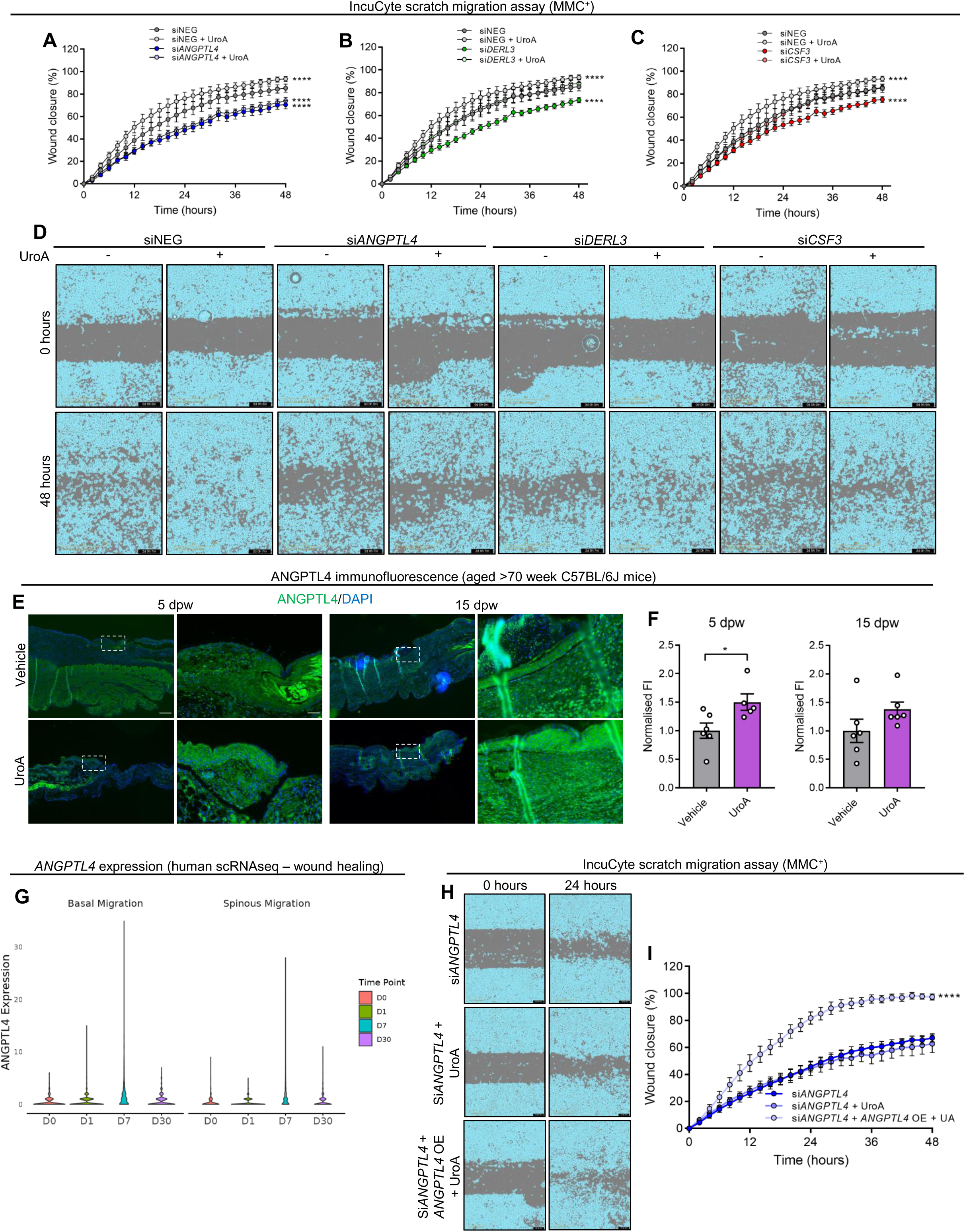
UroA promotes *ANGPTL4* expression to drive keratinocyte migration. (**A-C**) Quantification (mean ± SEM) of wound closure percentage in siNEG and (**A**) si*ANGPTL4,* (**B**) si*DERL3*, and (**C**) si*CSF3* HEKa treated with vehicle or UroA. Two-way ANOVA, **** = p < 0.0001. Each dot represents the mean of an individual biological replicate (n =3) containing at least 3 technical repeats. (**D**) Representative images of scratch wound migration at 0 and 48 hours. (**E**) Representative fluorescence images of ANGPTL4 staining in wound biopsies from aged mice at 5 days post wounding (dpw) and 15 dpw. Scale bar = 200μm (full size), 50μm (zoom). (**F**) Quantification (mean ± SEM) of ANGPTL4 fluorescence intensity (FI) normalised to vehicle in the leading edge epidermis in aged mice treated with vehicle or UroA. Students ttest, * = p < 0.05. Each dot represents an individual mouse (n = 11). (**G**) Violin plot of *ANGPTL4* expression in basal migration and spinous migration keratinocytes from scRNAseq analysis human wound healing. (**H**-**I**) Representative images and (**I**) quantification (mean ± SEM) of UroA rescue scratch wound assay in si*ANGPTL4* HEKa. Two-way ANOVA, **** = p < 0.0001. Each dot represents the mean of an individual biological replicate (n = 3) containing at least 3 technical repeats.

### UroA promotes FOSL1 transcription factor activity to enhance *ANGPTL4* transcription

We next sought to determine how UroA treatment led to the transcriptional upregulation of *ANGPTL4* in HEKa (**Figure 6A**). Firstly, enrichment analysis of transcription factor (TF) coexpression in significantly upregulated genes in UroA-treated HEKa from the RNAseq data identified the top TF to be FOSL1 (**Figure 6B**). Next, as FOSL1 is part of the AP-1 TF family, we then assessed the activity of several AP-1 TFs in UroA and vehicle-treated HEKa. Here, the activity of FOSB, FOSL1, and c-JUN was significantly increased in UroA-treated HEKa compared to vehicle (**Figure 6C**), with FOSL1 activity being the most significantly greater. There was no significant difference in c-FOS, JUND, or JUNB activity between UroA and vehicle-treated HEKa (**Figure 6E**). Immunofluorescence assessment additionally confirmed a significant increase in nuclear-localised FOSL1 staining in UroA-treated HEKa compared to vehicle (**Figure 6D**). Finally, *in sillico* analysis using JASPAR motif analysis confirmed the presence of 12 separate putative FOSL1-binding sites located within the *ANGPTL4* promoter region (**Figures 6F-G**). Collectively, these studies indicated that the UroA-mediated increase in *ANGPTL4* transcription was facilitated through increased FOSL1 TF activity.

**Figure 6.**
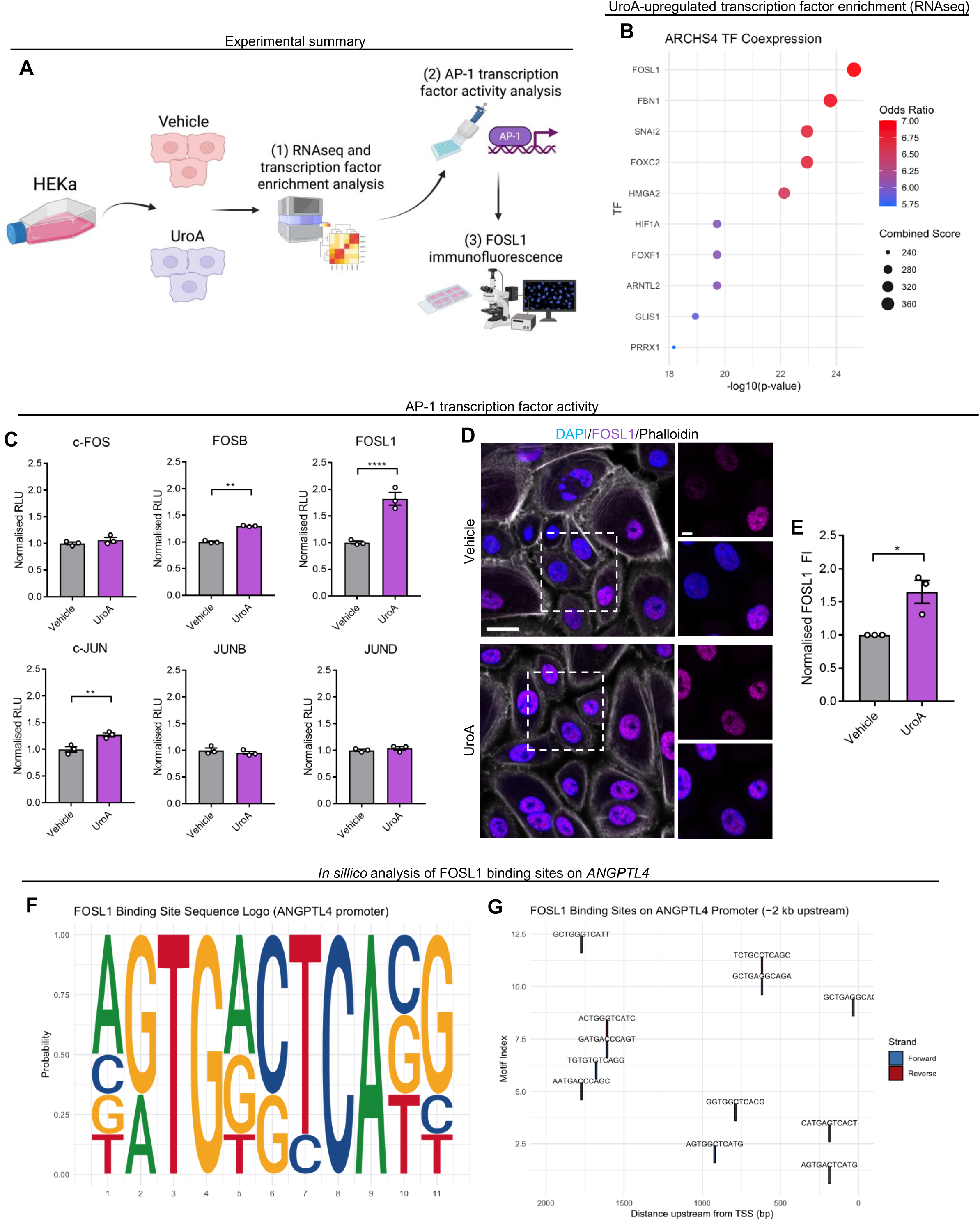
UroA promotes FOSL1 transcription factor activity to enhance *ANGPTL4* transcription. (**A**) Schematic diagram of the experimental plan. (**B**) Enrichment analysis showing ARCHS4 TF Coexpression of genes significantly upregulated in UroA-treated HEKa from RNAseq. (**C**) Quantification (mean ± SEM) of AP-1 family transcription factor activity in vehicle and UroA-treated HEKa. Students t-test, **** = p < 0.0001, ** = p < 0.01. Each dot represents the mean of an individual biological replicate (n = 3) containing at least 3 technical repeats. (**D-E**) Representative fluorescence images and (**E**) quantification (mean ± SEM) of FOSL1 staining in vehicle and UroA-treated HEKa. Scale bar = (full size), (zoom). Students t-test, * = p < 0.05. Each dot represents the mean of an individual biological replicate (n = 3) containing at least 3 technical repeats. (**F**) Motif analysis of putative FOSL1 binding sites in the *ANGPTL4* promoter region. (**G**) Genomics viewer of putative FOSL1 binding sites in the *ANGPTL4* promoter region. Binding sites are plotted by their motif index and distance upstream from the transcription starting site (TSS).

### UroA-induced metabolic reprogramming leads to AKT-GSK3 pathway activation

Following the demonstration that UroA led to increased FOSL1 TF activity, we sought to deduce the mechanism by which this occurred through UroA-mediated mitophagy. As mitophagy is known to induce alterations in cellular metabolism as well as reactive oxygen species (ROS) signalling^23, 24^, we utilised treatments with 2-dedoxyglucose (2-DG), an inhibitor of glycolysis; oligomycin, the inhibitor of complex II of the electron transport chain (ETC); and N-Acetyl-L-cysteine (NAC), a ROS scavenger. Firstly, cell viability assessment determined there to be no significant effect of any 24 hour treatment on HEKa cell viability (**Supplementary Figure 6A**). Next, RT-qPCR investigation of *ANGPTL4* gene expression following treatment showed that treatment with 2-DG significantly decreased *ANGPTL4* gene expression, whilst both oligomycin and NAC treatment significantly increased *ANGPTL4* gene expression in HEKa cells (**Figure 7A**). Interestingly, although UroA did not rescue the decreased *ANGPTL4* expression following 2-DG treatment, it did significantly decrease the elevated *ANGPTL4* expression induced by either oligomycin or NAC treatment (**Figure 7A**). This suggested that increases in glycolytic as opposed to oxidative metabolism, as well as decreases in ROS abundance led to elevated *ANGPTL4* gene expression.

**Figure 7.**
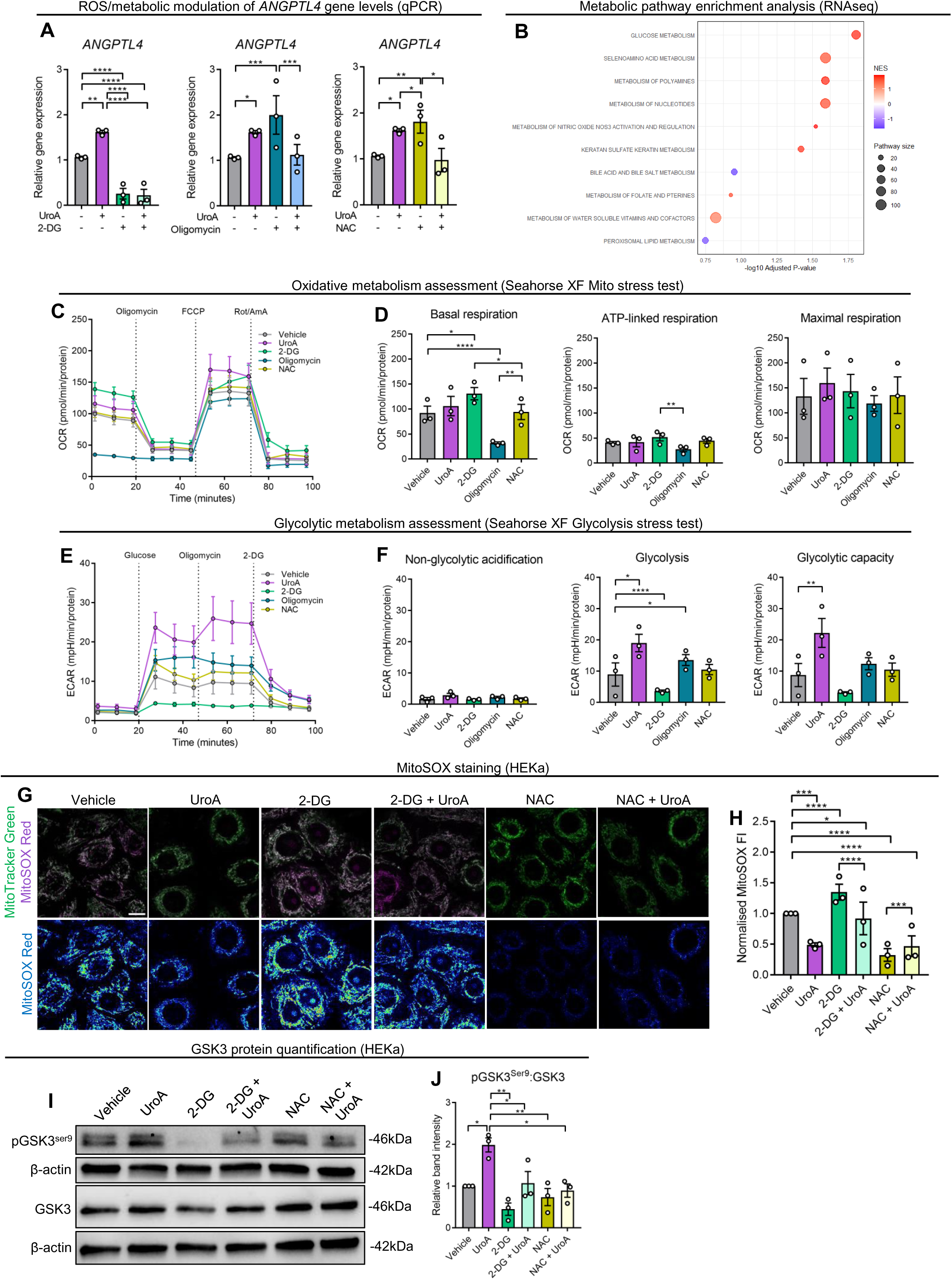
Upregulation in glycolytic metabolism and AKT-GSK3 pathway activity in UroA treated HEKa. (**A**) Quantification (mean ± SEM) of *ANGPTL4* gene expression in response to 2-DG, oligomycin, and NAC treatment in HEKa cells. Two-way ANOVA. Each dot represents the mean of an individual biological replicate (n = 3) containing at least 3 technical repeats. (**B**) Enrichment analysis of Reactome metabolic pathways upregulated in UroA-treated HEKa from RNAseq analysis. (**C**) Quantification (mean ± SEM) of OCR in Mito Stress Test assessments. Dots represent the mean of 3 biological replicates, each containing at least 3 technical replicates. (**D**) Quantification (mean ± SEM) of basal respiration, ATP-linked respiration, and maximal respiration in HEKa. Two-way ANOVA, **** = p < 0.0001, ** = p < 0.01, * = p < 0.05. Each dot represents the mean of an individual biological replicate (n = 3) containing at least 3 technical repeats. (**E**) Quantification (mean ± SEM) of ECAR in Glyco Stress Test assessments. Each dot represents the mean of an individual biological replicate (n = 3) containing at least 3 technical repeats. (**F**) Quantification (mean ± SEM) of non-glycolytic acidification, glycolysis, and glycolytic capacity in HEKa. Two-way ANOVA, **** = p < 0.0001, ** = p < 0.01, * = p < 0.05. Each dot represents the mean of an individual biological replicate (n = 3) containing at least 3 technical repeats. (**G-H**) Representative fluorescence images and (**H**) quantification (mean ± SEM) of MitoTracker Green and MitoSOX Red live cell staining in HEKa. **** = p < 0.0001, *** = p < 0.001, * = p < 0.05. Each dot represents the mean of an individual biological replicate (n = 3) containing at 15 cells. (**I-J**) Representative immunoblot and (**J**) quantification (mean ± SEM) of pGSK3 and GSK3 protein levels in HEKa. Two-way ANOVA, ** = p < 0.01, * = p < 0.05. Each dot represents the mean of an individual biological replicate (n = 3).

Next, to elucidate whether UroA treatment induced alterations in either of these factors, we firstly examined cellular metabolism in HEKa cells. Metabolic pathway enrichment analysis from the RNAseq demonstrated an upregulation in glycolytic metabolism as well as oxidative stress defence and metabolism of polyamines and nucleotides (**Figure 7B, Supplementary Figure 6B**). Next, seahorse assessment displayed no significant difference in the oxygen consumption rate (OCR) between UroA- and vehicle-treated HEKa, whilst 2-DG treatment significantly increased, and oligomycin significantly decreased basal respiration compared to vehicle (**Figure 7C-D**). When examining glycolytic metabolism (**Figure 7E**), both glycolysis and glycolytic capacity was significantly increased in UroA treated HEKa compared to vehicle, whilst glycolysis was significantly decreased in 2-DG treated, and significantly increased in oligomycin-treated HEKa compared to vehicle, as expected (**Figure 7F**). There was no significant difference in either oxidative or glycolytic metabolism in NAC-treated HEKa compared to vehicle. We then assessed mitochondrial ROS (mtROS) through MitoSOX Red live cell imaging of HEKa in similar conditions (**Figure 7G, Supplementary Figure 6C**). Here, both NAC and UroA treatment significantly decreased mtROS levels compared to vehicle, whilst treatment with 2-DG significantly increased mtROS levels (**Figure 7H**). Interestingly, additional treatment with UroA significantly decreased mtROS in 2-DG treated cells, and significantly increased mtROS in NAC-treated HEKa (**Figure 7H**). Overall, this suggested that UroA treatment led to an upregulation in glycolytic metabolism and a decrease in mtROS levels in HEKa cells. Finally, as the AKT-GSK3 signalling pathway is known to lead to increased FOSL1 TF activity and be upregulated by glycolysis and not primarily mediated through ROS signalling^25^, we next assessed whether UroA-mediated upregulation of glycolysis and decrease in ROS induced its pathway activation. Here, western blot for phosphorylated GSK3 (at serine 9) (pGSK3) demonstrated that UroA treatment significantly increased pGSK3:GSK3 ratio compared to vehicle, 2-DG, and NAC treated HEKa (**Figure 7I-J**).

### ANGPTL4 promotes laminin-332 dynamics in UroA-treated HEKa

After demonstrating how metabolic reprogramming induced by UroA resulted in elevated AKT-GSK3 pathway signalling to increase FOSL1-mediated *ANGPTL4* transcription, we next wanted to determine how ANGPTL4 promoted HEKa migration. Initially, as RNAseq analysis previously highlighted an elevation in genes associated with ‘laminin interactions’ and ‘nonmembrane ECM interactions’, as well as the fact that gene expression of several integrin and laminin-related genes was significantly increased in UroA-treated HEKa compared to vehicle (**Figure 8A**), we further investigated whether UroA-induced increase of these migration-mediating complexes was driven by ANGPTL4. Here, RT-qPCR analysis of *ITGA1, ITGB1* (integrin A2β1), *ITGA6, ITGB4* (integrin A6β4), *LAMA3,* and *LAMC2* (laminin-332) demonstrated that KD of *ANGPTL4* decreased the expression of both *LAMA3* and *LAMC2*, and was not rescued by UroA treatment (**Figure 8B**). Additionally, *ANGPTL4* OE increased the expression of these two laminin markers when compared to pCMV (**Figure 8C**). The expression of *ITGA1, LAMA1,* and *LAMB1* was significantly elevated in UroA treated siNEG compared to si*ANGPTL4* (**Supplementary Figure 7A**) further emphasising the importance of *ANGPTL4* and its downstream targets in UroA mediated increased keratinocyte migration, although not with *ANGPTL4* OE (**Supplementary Figure 7B**). To confirm this at the protein level, we next undertook immunofluorescence staining of laminin332 and phalloidin. Here, both whole cell laminin-332 and laminin-332 co-localised to the phalloidin cell boundary was significantly greater in UroA-treated siNEG cells compared to untreated siNEG, as well as both UroA treated and untreated si*ANGPTL4* cells (**Figure 8EF**). Finally, *ANGTPL4* KD significantly reduced whole cell and phalloidin boundary laminin322 when compared to siNEG, and was not rescued by UroA treatment. In addition, OE of *ANGPTL4* significantly increased whole cell and phalloidin boundary laminin-332 levels (**Figure 8G-H**). There were no differences in the distance from the cell nuclei and leading edge. Overall, these results suggested that UroA-mediated ANGPTL4 increases laminin-332 production which may be relevant for the physical aspects of cell migration (**Figure 8I**).

**Figure 8.**
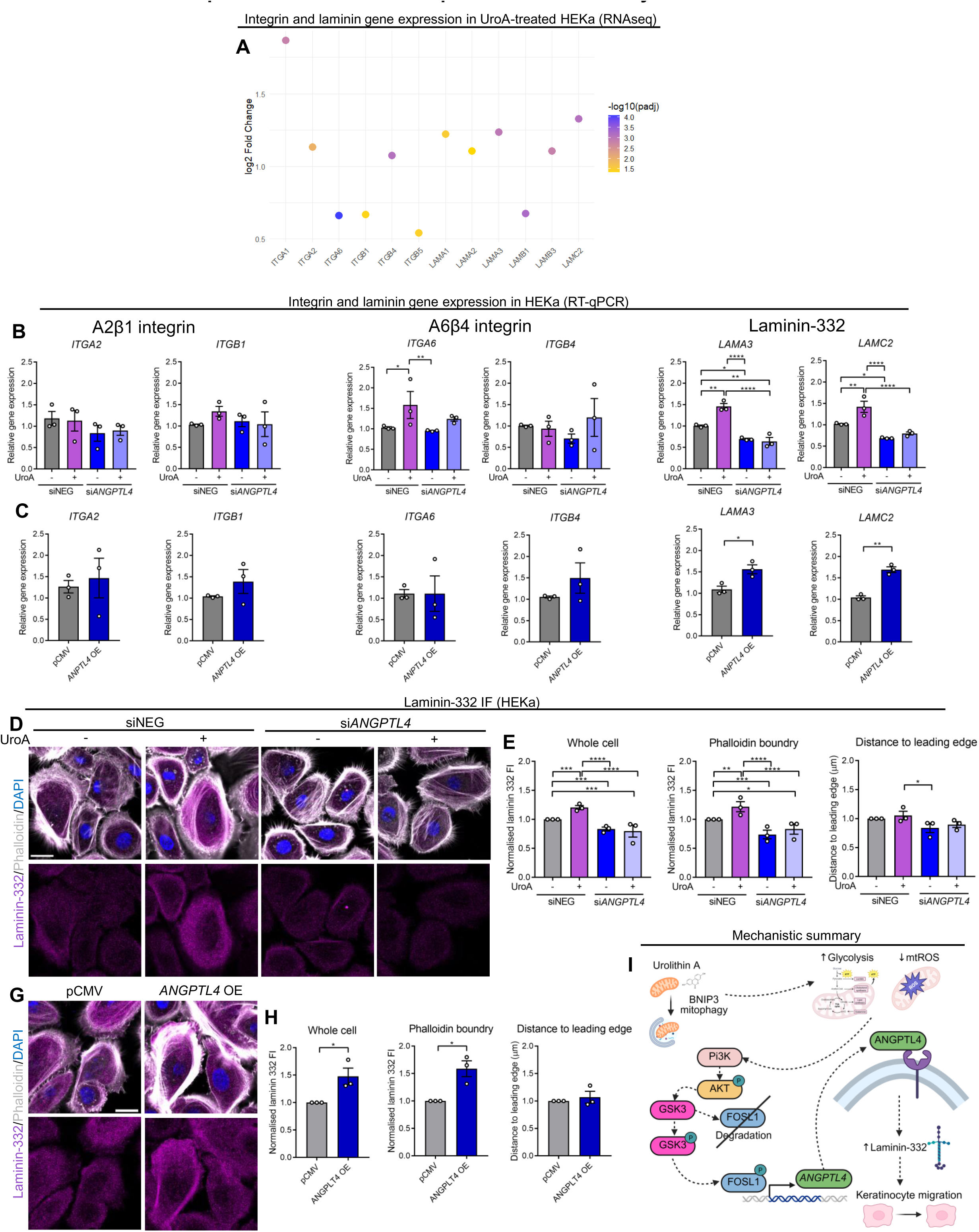
*ANGPTL4* promotes laminin-332 expression in keratinocytes. (**A**) Bubble plot depicting the Log2 fold change and −log10(padj) of integrin and laminin genes significantly upregulated in UroA-treated HEKa from RNAseq analysis. (**B**) RT-qPCR quantification (mean ± SEM) of integrin and laminin gene expression in transfected HEKa. Two-way ANOVA, **** = p < 0.0001, *** = p < 0.001, ** = p < 0.01, * = p < 0.05. Each dot represents the mean of an individual biological replicate (n = 3) containing at least 3 technical repeats. (**C**) Representative fluorescence images of HEKa stained with laminin-332, phalloidin, and DAPI. Scale bar = 20µm. (**D-H**) Quantification of normalised whole cell and phalloidin co-locailsed laminin-332 fluorescence intensity, as well as distance (µm) from the nuclei to leading edge. Two-way ANOVA, **** = p < 0.0001, *** = p < 0.001, ** = p < 0.01, * = p < 0.05. Each dot represents the mean of an individual biological replicate (n = 3) containing at 15 cells. (**H**) Schematic representation of the mechanistic summary.

## Discussion

A better understanding of the cellular and molecular mechanisms which underpin keratinocyte migration and thus re-epithelialisation during wound healing are essential in order to develop strategies to promote functional wound closure^5^. Utilising scRNAseq data of human wound healing, we firstly demonstrated that ub-independant mitophagy, particularly BNIP3-mediated mitophagy, is upregulated in migrating keratinocytes during the proliferation and re-epithelialisation stage of human wound healing. Gene co-regulation analysis suggested that increased *BNIP3* expression in migrating keratinocytes at day 7 was associated with metabolic pathways as well as pathways associated with gap junction and motility. This supports previous findings that BNIP3 mitophagy promoted the degradation of p-MAP4 to enhance keratinocyte migration and proliferation during wound healing^16^, as well as accelerating keratinocyte migration in non-wound healing migration contexts^17, 18^. *In vitro* studies confirmed that mitophagy was elevated in migrating keratinocytes, and that KD of *BNIP3* impaired keratinocyte migration, whilst induction of mitophagy through UroA increased migration. Additionally, treatment with UroA enhanced regeneration and genetic expression of important keratinocyte markers in zebrafish caudal fin amputation studies. Collectively, this is in concert with studies showing that mitophagy increases migration in contexts such as cancer cells^26, 27^.

In order to investigate the molecular mechanism by which mitophagy increased keratinocyte migration, we undertook RNAseq analysis of UroA-treated HEKa, which demonstrated an upregulation in genes involved in cell migration and ECM-related pathways, among others. To examine the impact of the most significantly upregulated genes on keratinocyte migration, we then performed a scratch wound migration screen after KD of the top 20 genes, which demonstrated *ANGPTL4, DERL3,* and *CSF3* to be the most important. Subsequent scratch wound migration experiments demonstrated that treatment of KD cells with UroA rescued migration in si*DERL3* and si*CSF3*, but not in si*ANGPTL4* HEKa, indicating that the migration enhancing effects of UroA are not mediated through DERL3 and CSF3, unlike with ANGPTL4.

ANGPTL4 is a member of the angiopoietin-like gene family involved in angiogenesis^28^, lipid metabolism^29^, and stem cell regulation^30^ and has been shown to play a beneficial role in wound healing promoting both keratinocyte migration and differentiation^31, 32^, reepithelialisation^32^, epithelial stem cell function^33^, macrophage differentiation^34^, as well as angiogenesis^35^. In this study, examination of *ANGPTL4* expression from scRNAseq data demonstrated an upregulation in migrating keratinocytes at the proliferation stage of wound healing, supporting previous studies which showed that ANGPTL4 is upregulated at the epidermis leading edge during wound healing^32, 36^. Importantly, immunofluorescence staining of wounded mice tissue performed in this study showed that ANGPTL4 was elevated at the epidermis leading edge in UroA-treated mice 5 dpw.

We next sought to investigate the upstream transcriptional regulators of *ANGPTL4* in UroA-treated HEKa. TF enrichment analysis of RNAseq data predicted several candidate TFs that were upregulated by UroA treatment, namely FOSL1 and AP-1 family TFs, and assessment of AP-1 TF activity demonstrated a significant upregulation of FOSL1 activity following UroA treatment. FOSL1 has a well-established role in regulating genes involved in cell migration and ECM remodelling^37^, including during wound healing^38^ and a recent study demonstrated the essential role of FOSL1 as a driver of re-epithelialisation^39^. Whilst FOSL1 and AP-1 TF family members are known to promote the transcription of genes involved in cell migration^40^, there is no previous demonstration that FOSL1 directly upregulates *ANGPTL4* transcription. To this end, *in sillico* analysis confirmed the presence of numerous putative FOSL1-binding sites on the *ANGPTL4* promoter region, reinforcing their link.

Mitophagy is known to promote metabolic reprogramming as well as modulate ROS signalling, which can collectively regulate redox-sensitive signalling pathways^41^, whilst ANGPTL4 is associated with lipid metabolism^29^. As such, we posited that UroA-mediated mitophagy altered HEKa cell metabolism and mitochondrial ROS levels to modulate redoxsensitive pathways and increase FOSL1 TF activity and *ANGPTL4* transcription. Herein, metabolic enrichment and Seahorse XF analysis demonstrated that UroA upregulated glycolysis and glycolytic capacity, whilst MitoSOX Red staining demonstrated a reduction in mitochondrial ROS. With regards to redox-sensitive signalling pathways, Pi3K-AKT-GSK3 pathway activation is known to be increased in energetic conditions as well as being redox-sensitive^25, 42^, and our results demonstrated the increase in phosphorylated GSK3 with UroA treatment, and a decrease with 2-DG treatment, supporting previous literature.

Phosphorylation and inhibition of GSK3 is the direct downstream target of AKT in the Pi3K-AKT pathway, which facilitates the stabilisation and nuclear accumulation of AP-1 family TFs^43^. Herein, our results demonstrate that an increase in glycolytic metabolism and decrease in mitochondrial ROS through UroA treatment led to Pi3K-AKT-GSK3 pathway activation and ultimately FOSL1 TF activity.

RNAseq analysis demonstrated a significant upregulation in various integrin and laminin genes, including those which make up the Α2β1 and Α6β4 integrins, as well as laminin-332, in UroA treated HEKa cells. ANGPTL4 has previously been shown to enhance keratinocyte migration through promoting interactions with β1 and β5 integrins to activate the FAK-srcPAK1 pathway^32^. Gene expression analysis of the aforementioned integrins and laminin-332 subunits in *ANGPTL4* KD and OE HEKa demonstrated that *ANGPTL4* KD significantly decreased levels of *LAMA3* and *LAMC2,* and was not rescued by additional UroA treatment, whilst expression was significantly greater after *ANGPTL4* OE – suggesting that UroAmediated upregulation in ANGPTL4 increased laminin-332 in HEKa. Laminin-332 is a large extracellular glycoprotein produced by keratinocytes and plays essential roles in cellular and ECM interactions^44^. As well as being a major component of anchoring filaments and interactions with integrins Α2β1 and Α6β4 to promote adhesion^45^, laminin-332 is involved in the formation of protrusive filopodial structures to enhance keratinocyte migration^46, 47^. Additionally, increased transcription and deposition of laminin-332 by leading edge keratinocytes is an important mediator of keratinocyte migration during wound healing, allowing for the interaction and activation of matrix metalloproteinases (MMPs) and ECM components^48, 49^. We showed through IF staining of laminin-332 and phalloidin in *ANGPTL4* KD and OE HEKa that UroA-mediated ANGPTL4 was important in both whole cell and cell edge accumulation of laminin-332.

In conclusion, we have shown in this study that mitophagy is upregulated in migrating keratinocytes at the proliferation stage of wound healing. By promoting mitophagy with UroA, we demonstrate that mitophagy induction enhanced regeneration in zebrafish caudal fin models, as well as accelerated keratinocyte migration *in vitro*. Through promoting metabolic reprogramming and the glycolytic capacity of keratinocytes, UroA-induced mitophagy increased AKT-GSK3 pathway and FOSL1 transcription factor activity to enhance *ANGPTL4* transcription and laminin-332 production. Collectively, these findings build upon previous studies to reinforce the idea that mitophagy is important during wound healing, uncover a previously unknown link between mitophagy and metabolism in wound healing contexts, and demonstrate for the first time an important role for mitophagy in AKT-GSK3 pathway activity and laminin-332 mediated migration of keratinocytes during wound healing. In doing so, this study indicates the potential benefit of targeting mitochondrial-related functions therapeutically in the wound healing context.

### Methods and materials

#### Ethics

This study was approved by Stockholm Regional Ethics Committee and executed in agreement with the Helsinki Declaration. Informed consent was obtained from all the research subjects.

#### Single-cell RNA sequencing data processing and keratinocyte subtype annotation

Previously reported^39^ raw scRNA-seq data from human wound healing skin samples was processed using the Seurat (v4.x) R package. Quality control filtering was applied to remove cells with low gene counts, high mitochondrial gene expression, or other quality metrics indicating poor data quality. The data were normalized and integrated across samples using SCTransform normalization and Harmony batch correction to mitigate batch effects between patients and time points.

Cells were clustered using shared nearest neighbour (SNN) modularity optimization clustering and visualized with Uniform Manifold Approximation and Projection (UMAP). Based on canonical marker genes (e.g. *KRT14, KRT5* for basal, *KRT1, IVL* for spinous), nine keratinocyte subtypes were manually annotated consistent with previously published classifications.

#### Pseudobulk differential expression analysis across wound healing time points

To investigate temporal gene expression changes within keratinocyte subtypes, a pseudobulk approach was used. For each keratinocyte subtype, raw counts were summed across cells belonging to the same patient and wound healing time point (D0, D1, D7, D30), generating sample-level gene count matrices. Differential expression (DE) analysis was performed using the limma-voom pipeline. The design matrix included time point as a fixed effect, and patient identity was modelled as a blocking factor using the duplicateCorrelation function to account for repeated measures. Six pairwise contrasts were tested to capture gene expression changes between wound healing stages: D1 vs D0, D7 vs D0, D30 vs D0, D7 vs D1, D30 vs D1, and D30 vs D7.

Lowly expressed genes were filtered using the filterByExpr function from edgeR, and count data were normalized using the trimmed mean of M-values (TMM) method. Empirical Bayes moderated t-statistics were calculated to identify differentially expressed genes at an adjusted p-value (FDR) threshold of 0.05.

#### Mitochondrial gene subset and visualization

Mitochondrial-related genes were extracted based on the MitoCarta 3.0 gene list, which catalogues human genes with mitochondrial localisation or function. The DE results were subset to this list to focus on mitochondrial dynamics during wound healing.

Volcano plots and heatmaps were generated for mitochondrial genes across keratinocyte subtypes and time points to visualize fold changes and significance. Heatmaps were clustered by gene and condition, displaying log2 fold change values to identify patterns of mitochondrial gene regulation.

#### Cell culture and treatments

HEKa were purchased from (Thermo Fisher Scientific) and cultured in EpiLife serum free keratinocyte growth media (Thermo Fisher Scientific) supplemented with human keratinocyte growth supplement (HKGS) (Thermo Fisher Scientific) and antibiotics (100 units/mL penicillin and 100 μg/mL streptomycin) (Thermo Fisher Scientific) in an incubator maintained at 37°C and 5% CO_2_. HEKa were treated with 5μM UroA (Sigma Aldrich), for 24 hours. For metabolism and ROS experiments, HEKa were treated with either 50mM 2-DG (Sigma Aldrich), 1μM oligomycin (Sigma Aldrich), or 5μM NAC (Thermo Fisher Scientific) for 24 hours.

#### Transfections

*BNIP3* KD was performed in HEKa using 25μM ON-TARGETplus human siRNA SMARTpool (Horizon Discovery) targeted to *BNIP3* (si*BNIP3*) for 24 hours with Lipofectamine RNAiMax (Thermo Fisher Scientific). In the scratch migration assay and RT-qPCR fishing experiments, KD of *FAM196B* (si*FAM196B*), *CXCL5* (si*CXCL5*), *CSF3* (si*CSF3*), *SPDEF* (si*SPDEF*), *ANGPTL4* (si*ANGPTL4*), *DERL3* (si*DERL3*), *COL8A1* (si*COL8A1*), *PLAT* (si*PLAT*), *MRGPRX4* (si*MRGPRX4*), *MRGPRX3* (si*MRGPRX3*), *TMEM106A* (si*TMEM106A*), *PPP1R1B* (si*PPP1R1B*), *IL1RL1* (si*IL1RL1*), *NTM* (si*NTM*), *CDH4* (si*CDH4*), *ADAMTSL1* (si*ADAMTSL1*), *MYCBPAP* (si*MYCBPAP*), *IL13RA2* (si*IL13RA2*), *GNG11* (si*GNG11*), and *DAW1* (si*DAW1*) was performed in HEKa using 25μM ON-TARGETplus human siRNA SMARTpool targeted to the respective gene for 48 hours using Lipofectamine RNAiMax. KD of *ANGPTL4*, *DERL3*, and *CSF3* in subsequent experiments was performed for 6 hours in serum-free OMTI-MEM (Thermo Fisher Scientific). Simultaneously, HEKa were transfected with universal negative control siRNA (siTOOLS Biotech) for similar durations to generate siNEG HEKa. *ANGPTL4* OE was was induced by *ANGPTL4* human-tagged ORF clone (NM_001039667) (Origene), with pCMV6 entry vector (Origene) acting as a nontargeting control, both for 6 hours using Lipofectamine 3000 (Thermo Fisher Scientific) in serum-free OMTI-MEM media. For double transfection experiments, HEKa were initially transfected with siRNA targeting *ANGPTL4* for 6 hours using Lipofectamine RNAiMax before overnight incubation in EpiLife media and subsequent transfection with *ANGPTL4* human-tagged ORF clone for 6 hours using Lipofectamine 3000 the next day. Transfection with universal negative control siRNA and pCMV6 entry vector was undertaken to generate negative controls.

#### Cellular viability

The cell viability at various concentrations of all pharmacological metabolic drugs was determined using resazurin-based PrestoBlue (Thermo Fisher Scientific). After incubation with PrestoBlue for 30 minutes, cell absorbance at 560nm and 600nm was monitored using a SpectraMax iD3 plate reader (SpectraMax) and the absorbance values were used to analyse cellular viability. HEKa were pre-treated for 2 hours with 3% hydrogen peroxide (H_2_O_2_) (Thermo Fisher Scientific) as a positive control.

#### Protein extraction and western blotting

Whole cell lysate fractions were isolated from HEKa as follows: cell pellet obtained was resuspended in RIPA buffer (Thermo Fisher Scientific) and kept at ice for 30 minutes with the inversion at 10-minute intervals. Resuspended cells then were centrifuged at 13000g for 10 minutes and the cell supernatant was subsequently used for western blotting. Isolated subcellular fractions (30-50ng) were loaded on 4-20% gradient SDS polyacrylamide gels (Biorad) and ran at 80V followed by 120V for 60-90 minutes. Gel was transferred onto nitrocellulose membrane and blocked in 5% skim milk (Thermo Fisher Scientific) or 5% bovine serum albumin (Thermo Fisher Scientific) diluted in TBST (Tris Buffered Saline with 0.05% Tween20) followed by incubation with primary antibody at 4°C overnight with slow shaking. On the next day, membranes were washed with TBST three times for ten minutes, incubated with HRP-conjugated secondary antibody for 1 hour, washed with TBST, and detected with femto ECL solution (Thermo Fisher Scientific). The Anti-rabbit or Anti-mouse secondary antibody was used at 1:5000 dilution. Information about antibodies used in these and other experiments are described in **Table 1**.

#### Mitophagy flux western blot

Protein levels of BNIP3 (Cell Signaling Technology) in WT HEKa in a mitophagy flux assay. In order to inhibit the degradation of lysosomal contents and recycling of mitophagy receptors, HEKa were treated acutely with 25μM CQ (Sigma Aldrich) for 1 hour prior to lysate harvesting and protein extraction as detailed above. For analysis of BNIP3 protein levels in siNEG and si*BNIP3* HEKa, cells were treated with 1μM DFP (Santa Cruz Biotechnology) for 24 hours before lysate harvesting and protein extraction. Western blot quantification of proteins was performed as described above.

#### RT-qPCR

Total RNA was extracted from cells using Trizol reagent (Thermo Fisher Scientific). After determining the sample quality, reverse transcription was performed using the RevertAid First Strand cDNA Synthesis Kit (Thermo Fisher Scientific). Gene expression was quantified by SYBR Green expression assays (Thermo Fisher Scientific) and normalized with β-actin. The primer information used in these experiments are listed in **Table 2**.

#### RNAseq gene expression analysis of UroA-treated HEKa

HEKa treated with either vehicle or 5μM UroA (n = 4 biological repeats per condition) underwent RNA extraction as stated above and RNA quality and quantity was analyzed with Nanodrop 1000, Qubit 4.0 Fluorometer (Life Sciences), and Agilent 5300 Fragment Analyzer (Agilent). RNA from samples were then sent for gene expression profiling using Illumina NovaSeq with Poly(A) selection at the Azenta Life Sciences, Germany and RNA sequencing libraries created using NEBNext Ultra RNA Library Prep Kit for Illumina (NEB) following manufacturer’s instructions. Briefly, mRNAs were first enriched with Oligo(dT) beads. Enriched mRNAs were fragmented for 15 minutes at 94°C. First strand and second strand cDNAs were subsequently synthesised. cDNA fragments were end repaired and adenylated at 3’ends, and universal adapters were ligated to cDNA fragments, followed by index addition and library enrichment by limited-cycle PCR. Sequencing libraries were validated using NGS Kit on the Agilent 5300 Fragment Analyzer, and quantified by using Qubit 4.0 Fluorometer. The sequencing libraries were multiplexed and loaded on the flowcell on the Illumina NovaSeq X plus instrument according to manufacturer’s instructions. The samples were sequenced using a 2×150 Pair-End (PE) configuration v1.5. Image analysis and base calling were conducted by the NovaSeq Control Software v1.7 on the NovaSeq instrument. Raw sequence data (.bcl files) generated from Illumina NovaSeq was converted into fastq files and de-multiplexed using Illumina bcl2fastq program version 2.20. One mismatch was allowed for index sequence identification. Sequence reads were trimmed to remove possible adapter sequences and nucleotides with poor quality using Trimmomatic v.0.36. The trimmed reads were mapped to the Homo sapiens reference genome available on ENSEMBL using the STAR aligner v.2.5.2b. The STAR aligner is a splice aligner that detects splice junctions and incorporates them to help align the entire read sequences. BAM files were generated as a result of this step. Unique gene hit counts were calculated by using feature Counts from the Subread package v.1.5.2. Only unique reads that fell within exon regions were counted. A PCA analysis was performed using the "plotPCA" function within the DESeq2 R package. The plot shows the samples in a 2D plane spanned by their first two principal components. After extraction of gene hit counts, the gene hit counts table was used for downstream differential expression analysis. Using DESeq2, a comparison of gene expression between the groups of samples was performed. The Wald test was used to generate p-values and Log2 fold changes. Genes showing at least 0.5-fold change with pvalue <0.05 after treatment were used for creating a list of differentially expressed genes. Enrichr analysis (https://maayanlab.cloud/Enrichr/) ^50^ was performed to understand the functionally altered pathways in cells treated with UroA and CQ compared to vehicle.

#### *In sillico ANGPTL4* binding site analysis

Putative FOSL1 binding sites within the *ANGPTL4* promoter region were identified using the JASPAR database^51^. The canonical FOSL1 position weight matrix (MA0477.1) was used for scanning the *ANGPTL4* promoter sequence, which was retrieved for transcript ENST00000167772 from the Eukaryotic Promoter Database (EPD)^52^. Binding sites were identified using a relative profile score threshold of 80%. Predicted motifs were mapped relative to the *ANGPTL4* transcription start site and annotated by strand orientation. Visualization of the binding motifs index was performed using R (version 4.5.0, R Core Team, 2024). Putative binding site sequences were visualized using sequence logos created with the ggseqlogo package^53^ in R.

#### Live cell staining of mitophagy and mitochondrial morphology during *in vitro* migration

HEKa were seeded onto μ-dish 35mm with a 4 well insert (Ibidi). When cells were confluent, the inserts were removed using a sterilised tweezer, creating a 500μm gap between cell populations. In order to quantify mitophagy, mitochondrial mass, and mitochondrial morphology at different stages of wound healing, cells were stained with 100nM MitoTracker Green (MTGr), 100nM Tetramethylrhodamine (TMRE), and 60nM LysoTracker Deep Red (LTDR) (all Thermo Fisher Scientific) for 15 minutes. After staining, the dye was washed out with three rinses of Dulbecco’s Phosphate Buffered Saline (PBS) (Thermo Fisher Scientific). Finally, a 100nM TMRE imaging bath was added before the cells were imaged at 63x magnification.

#### MitoSOX Red live cell staining and imaging

HEKa were seeded into μ-slide eight-well glass bottom plates (Ibidi). Following treatment, cells were stained with NAC and 100µM hydrogen peroxide (H_2_O_2_) for 30 minutes to act as a positive and negative control respectively. HEKa were then stained with 500nM MitoSOX Red (Thermo Fisher Scientific) (containing respective treatments) for 30 minutes in an incubator maintained at 37°C and 5% CO_2_. After three rinses with Ca^2+^-free Hanks Buffered Saline Solution (HBSS), cells were incubated with 100nM MTGr for ten minutes. After rinsing with HBSS and addition of respective treatments, cells were imaged at 63x magnification.

#### Laminin-332 immunofluorescence

Immunofluorescence (IF) of Laminin-332 and phalloidin was performed on HEKa seeded into μ-slide eight-well glass bottom plates (Ibidi) according to manufacturers instructions. Briefly, following transfection and treatment, cells were fixed with 4% paraformaldehyde (PFA) (Thermo Fisher Scientific) for 10 minutes before washing with PBS (thermo Fisher Scientific) and permeabilisation with 0.1% TritonX-100 (Sigma Aldrich) for 10 minutes. Following, cells were washing with PBS and then blocked with 5% BSA for 1 hour at room temperature (RT) before incubation with primary Laminin-332 antibody (Thermo Fisher Scientific) for 1 hour at RT. After washing with PBS, cells with incubated with secondary antibody and Phalloidin i555 (1:1000) (Abcam) for 1 hour at RT. Finally, following washing with PBS, cells were incubated with DAPI (1:2000) for 5 minutes at RT, washed with PBS, and stored at 4^°^C until imaging.

#### FOSL1 immunofluorescence

IF of FOSL1 in HEKa was performed according to manufacturers instructions. Briefly, HEKa were seeded into μ-slide eight-well glass bottom plates. After treatment with UroA or vehicle for 24 hours, cells were fixed with 4% PFA for 10 minutes, permeabilised with 0.1% TritonX-100 for 10 minutes, and blocked with 5% BSA for 1 hour, before incubation with FOSL1 ab (1:500) (Thermo Fisher Scientific) for 1 hour at RT. Next, cells were incubated with secondary antibody for 1 hour at RT, followed by DAPI (1:2000) for 5 minutes at RT, washed with PBS, and stored at 4^°^C until imaging.

#### Confocal microscopy and image analysis

IF and live cell imaging was performed on a Zeiss LSM880 with airyscan. With regards to the quantification of mitochondrial factors from live cell imaging of HEKa, cells were classed into three categories depending on their proximity to the wound: stationary, which were localised in the main body of cells far away from the wound; wound edge (WE), which were found on the boundary of the wound; and migrating, which were in the gap between the two cell populations. For the quantification of mitophagy, the number of foci of colocalized MTGr, TMRE, and LTDR events was determined for each cell. The Mitochondria Analyzer ImageJ plugin^54^ was used to quantify mitochondrial mass, aspect ratio (AR), and form factor (FF). All images were acquired as Z-stacks at 63x magnification. Three biological repeats with 30 cells for each category were acquired at 0, 24, 48, 72, and 96 hours after removal of the insert.

For live cell imaging of MitoSOX Red and MTGr, cells were imaged using channels for Alexa 547 (MitoSOX Red) and Alexa 488 (MTGr), with channel settings determined using a noprimary control (NPC). Following acquisition at 63x magnification, fluorescence intensity was quantified and normalised to the fluorescence intensity of the mean vehicle control for each biological repeat using ImageJ software.

For imaging of laminin-332 and phalloidin-stained HEKa, cells were imaged using the Alexa 488 (laminin-332), Alexa 547 (phalloidin), and DAPI (nuclei) channels at 63x magnification, with laser strength and gain set following after normalization to NPC. All images were acquired as Z-stacks, with three biological repeats containing 15 cells in each condition. With regards to analysis, laminin-332 fluorescence intensity was quantified for both the whole cell, as well as area of the cell co-localised with phalloidin on ImageJ software. Measurements of the distance from the leading edge to nuclei was undertaken on Zen Blue (Zeiss).

For imaging and analysis of nuclear-localised FOSL1 in HEKa, cells were imaged using Alexa 547 (FOSL1) and DAPI channels at 63x magnification with laser strength and gain set following after normalization to NPC. For analysis, the fluorescence intensity of FOSL1 colocalised with the cell nucleus was quantified for 15 cells in three separate biological repeats using ImageJ software, with normalization to the mean FI of vehicle cells in the respective biological repeat.

#### Seahorse XF bioenergetic assays

HEKa cells were seeded on XFe24-well cell culture microplates (Agilent) at 2.8−3.5 × 10^4^ cells/well for 16−24 hours. For the Mito Stress Test, OCR (percentage oxygen consumption rate over baseline) was measured using an XFe24 extracellular flux analyser (Agilent) in a minimal DMEM assay medium supplemented with 10mM glucose (Thermo Fisher Scientific), 1mM sodium pyruvate (Thermo Fisher Scientific), and 2mM L-glutamine (Thermo Fisher Scientific) for 45−60 minutes for equilibration. Compound injections included 2µM oligomycin, 1µM FCCP, and 0.5µM rotenone + antimycin A (both Thermo Fisher Scientific). For the Glycolysis Stress Test, ECAR (percentage extracellular acidification rate over baseline) was measured in cells cultured in minimal DMEM assay medium supplemented with 2mM L-glutamine for 45−60 minutes for equilibration. Compound injections included 10mM glucose, 2µM oligomycin, and 50mM 2-DG. For both tests, measurements were taken before and after exposure to compound injections at regular intervals for 3 cycles. Data was analysed using the Wave Software (Agilent) and expressed as a percentage of oxygen consumption rate or extracellular acidification rate change over the pre-exposure baseline rate.

#### Aged zebrafish fish husbandry and ethics

Project authorisation was obtained from the Animal Experimental Board of the Regional State Administrative Agency for Southern Finland (licence numbers ESAVI/31414/2020 and ESAVI/29099/2024). Aged zebrafish (158-164 weeks) were maintained at the Zebrafish Core at Turku Bioscience Centre, University of Turku and Åbo Akademi University, Turku, Finland according to standard procedures. Zebrafish were fed with Gemma micro dry food.

Wild-type zebrafish (AB strain) embryos were exposed to various concentrations of UroA (BOC Sciences) (diluted in 1% DMSO) from 2 days post fertilisation (dpf) to 4 dpf in order to determine potential toxicity and behavioral effects. Initially, zebrafish embryos in 96 well glass-bottom plates were photographed using a Nikon Eclipse Ti2-Emicroscope with 2x Nikon Plan-Apochromat (NA0.06) objective and Hamamatsu sCMOS Orca Flash4.0 camera in order to assess morphological features. Heart rate imaging was performed with same instrument using time-lapse imaging with 50ms intervals and 2ms exposure using 2×2 binning in camera. Morphological features and heart rate were analysed using DanioScope software v1.1 (Noldust IT). Behavioral experiments were carried out using DanioVision instrument (Noldus IT, Netherlands). Embryos were exposed in square-well 96-well plates from 2dpf and motility analysed at 4dpf. Each well contained 1 embryo movement was tracked over the course of 70 minutes with 20 minutes of baseline and 5 cycles of alternating light pulses of 5min light and 5min darkness to increase embryo motility. Embryo tracking and movement analysis done using Ethovision XT 13 software (Noldus IT)

#### Aged zebrafish caudal fin regeneration assay

Wild-type aged zebrafish (AB strain) were used for studying the effect of UroA in aged zebrafish caudal fin regeneration model^55^. At day 0, fish were anesthetized with 200mg/l tricanine methanosulfonate (MS222,Sigma Aldrich) diluted in system water before being placed on a petri dish with a tablespoon and having their caudal spread on top of the lid. Following, uncut caudal fins were photographed using a Zeiss Stemi DV4 stereomicroscope and mobile phone before a small section of distal caudal fin was resected with a scalpel. Following subsequent imaging (day 0, cut) fish were allowed to recover from anesthesia and housed individually in a 1L tank containing either 10µM UroA (n =20) or DMSO (n=20) in 0.005% methylene blue and 5 mg/l lidocaine in system water and placed in a warm room (26°C) for the experiment duration. Water parameters were followed and tank water changed partially every 2 days and fully every 4 days. At days 4 and 8 post amputation, fish were again anesthetised and their caudal fins photographed under the stereomicroscope. At both days 0, 4, and 8 resected caudal fins were split in half, with half the section to be used for RT-qPCR analysis frozen and stored at −80°C and the other half fixed in 10% neutral buffered formalin. With regards to analysis, the caudal fin thickness of cut fins at day 0 was measured before the thickness of the regeneration zone (from the resection plane to the distal end of the regenerating fin) was measured at days 4 and 8 post amputation and quantified as a percentage of regeneration.

#### Aged zebrafish caudal fin RNA extraction and RT-qPCR

RNA from resected fin pieces was extracted using NucleoSpin RNA Plus XS kit for RNA purification (Machery-Nagel) according to manufacturers protocol and stored at −80°C. RTqPCR was performed using SYBR Green assay as described above. Primer information is described in **Table 2**.

#### Aged murine wound healing model histochemistry staining and imaging

In a separate study, aged female C57BL/6 mice (>70 weeks) (Janvier Labs, France) (n = 24) were administered daily with either 25mg/ml UroA (BOC Sciences) or vehicle (0.5% w/v carboxymethlycellulose, 0.25% v/v, Tween 80 in dH_2_O) through oral gavage and underwent wound healing assessment following the administration of 6mm full-thickness biopsies. Protocols were assessed and approved by Ethical Committee N°CEEA-122, and animals were housed at the animal facilities of the Zone d’Evaluation Fontionnelle (ZEF), France.

With regards to wounding, following a 5 day acclimitisation period, wounds were taken following shaving of the back under isoflurane anesthesia and immediately splinted with silicone rings and allowed to heal under moist conditions. At days 5 and 15 post wounding, wounds were harvested and either stored in 4% PFA or immediately snap-frozen and placed in dry ice. Following harvesting of wounds, animals were euthanised.

Following sectioning of formalin-fixed paraffin-embedded (FFPE) blocks (10µm thickness), 5 and 15 dpw biopsies from aged mouse wound models underwent immunofluorescence staining to assess ANGPTL4 expression at the wound leading edge, according to manufacturer instructions. Briefly, Slides were incubated at 65°C for 1 hour before deparafinisation and rehydration along an ethanol gradient as standard. Next, sections were washed in TBST three times for 5 minutes, followed by antigen retrieval in 10mM sodium citrate pH 6.0 for 7 minutes and blocking with 5% BSA for 1 hour at RT. Following a wash cycle with TBST, sections were incubated with ANGPTL4 (Thermo Fisher Scientific) primary antibody 1:500) for 1 hour at RT. After washing, sections were then incubated with secondary antibody for 1 hour at RT, washed, and then incubated with DAPI (1:2000) for 5 minutes at RT and finally stored at 4°C. For imaging and analysis of ANGPTL4 expression in the leading-edge epidermis, sections were imaged using a Nikon Eclipse NI-E upright fluorescent microscope at 4x and 20x magnification using the Cy3 channel. ANGPTL4 fluorescence intensity was quantified for each section and normalized to the mean of the vehicle-treated mice on ImageJ.

#### Scratch wound healing assay

HEKa were seeded on 96-well ImageLock plates (Sartorius) that were pre-coated with 300µg/ml rat collagen-I (Thermo Fisher Scientific) for 24 hours. After the cells reached monolayer confluence they were scratched with the Wound Maker 96 (Essen Bioscience) in order to generate wounds. In addition, where stated, cells were treated with and without a 2 hour pre-treatment with 10μg/ml mitomycin c (MMC) (Sigma Aldrich) prior to scratching, in order to inhibit cell proliferation. After scratching, the cells were incubated in an IncucyteZOOM (Essen Bioscience) and imaged every 2 hours at 10x magnification for 48-72 hours. Relative wound density (%) was quantified on Incucyte 2022A software (Sartorius).

#### AP-1 Transcription factor activity assay

The DNA binding activity of AP-1 family transcription factors (c-Fos, FosB, FOSL1, c-Jun, JunB, and JunD) in vehicle and UroA-treated HEKa was determined using the AP-1 Transcription Factor Assay Kit (Colorimetric) (Abcam, ab207196) according to manufacturer instructions. Briefly, nuclear extracts were prepared from HEKa using the Nuclear Extraction Kit (Abcam, ab113474) with protein concentrations determined by BCA assay. Equal amounts (2μg) of nuclear protein were incubated in 96 well plates coated with oligonucleotide containing the respective AP-1 binding sequence as well as competitor or mutant oligonucleotide controls before bound complexes were detected by incubation with primary antibody followed by anti-rabbit IgG HRP-conjugated secondary antibody. After washing to remove unbound proteins, TMB substrate developing solution was added before the reaction was stopped and absorbance measured at 450nm and 665nm on a SpectraMax iD3 plate reader. Background signal was subtracted and the values were normalised to vehicle.

#### Data availability and reproducibility

Raw RNAseq data is has been deposited in the NCBI Sequence Read Archive (SRA) with the accession numbers: SAMN52852917, SAMN52852918, SAMN52852919, SAMN52852920, SAMN52852921, SAMN52852922, SAMN52852923, SAMN52852924. The datasets generated and/or analysed during the current study are available from the corresponding authors on reasonable request.

#### Statistical Analysis

Statistical significance was determined by two tailed Student’s t-test. The significance among two group was determined by paired or unpaired t-tests, and by Two-way ANOVA for multiple groups, both on GraphPad Prism Version 6. The p-values of GSEA analysis were calculated by using Fisher’s exact test. P-value < 0.05 was determined to be statistically significant.

## Acknowledgments

We thank the patients for participating in this study and providing the biopsies as well as research nurse Helena Griehsel. This work was supported by grants from Hudfonden, Swedish Science Council, Swedish Society for Medical Research, Åke Wibergs Stiftelse, LEO foundation, ALF Medicin Stockholm, Jeanssons Stiftelse, Wallenberg foundation, Tore Nilssons Stiftelse, and Stiftelsen Sigurd and Elsa Goljes Minne.

## Author contributions

MH, JW, and EBW conceptualised the project and planned the experiments. MH, MT, NW, JV, IP, MC, GAQ, and EBW performed experiments and analysed data. All authors contributed to the manuscript writing and editing.

---

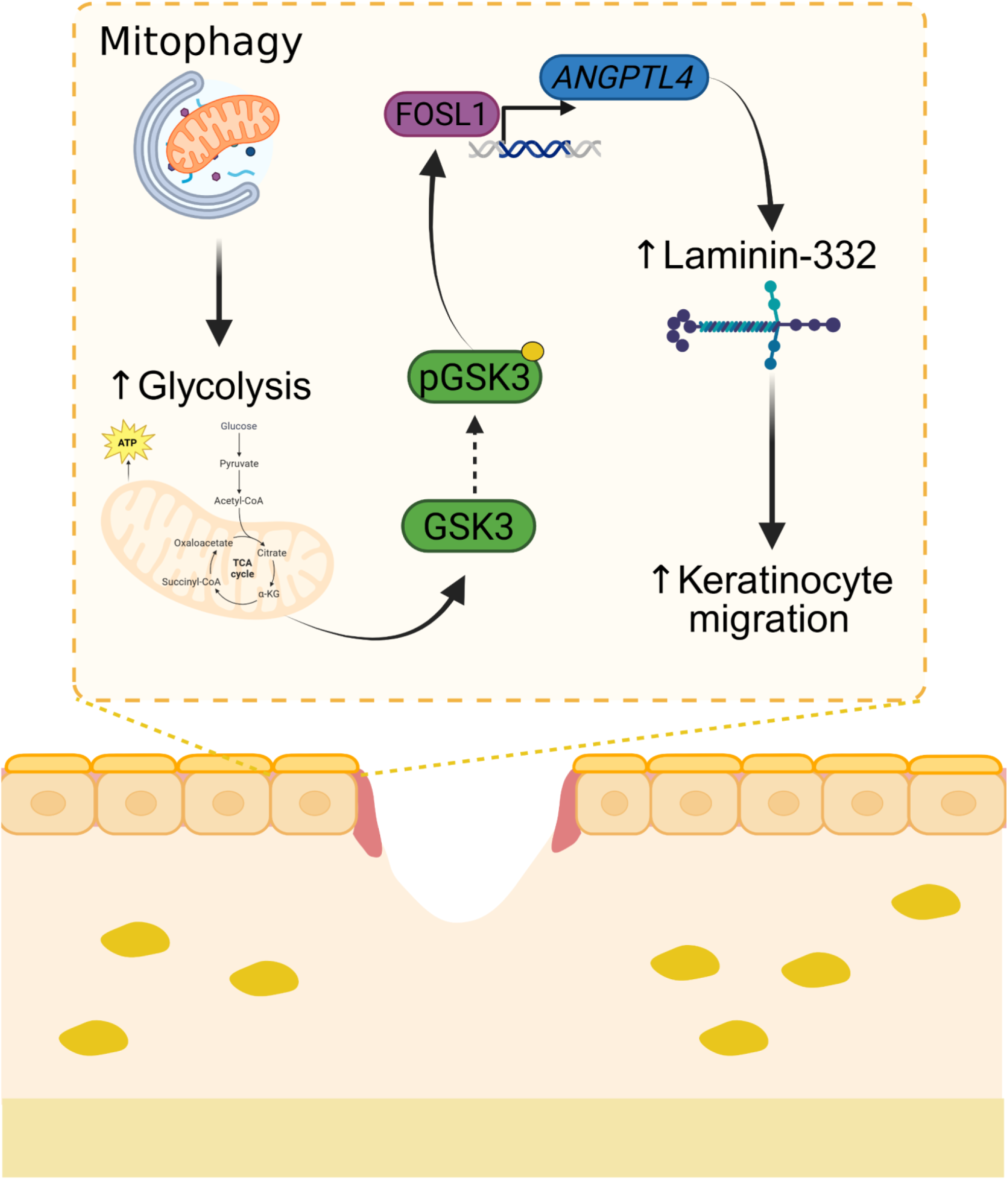

## Supplemental information

**Supplementary Figure 1.**
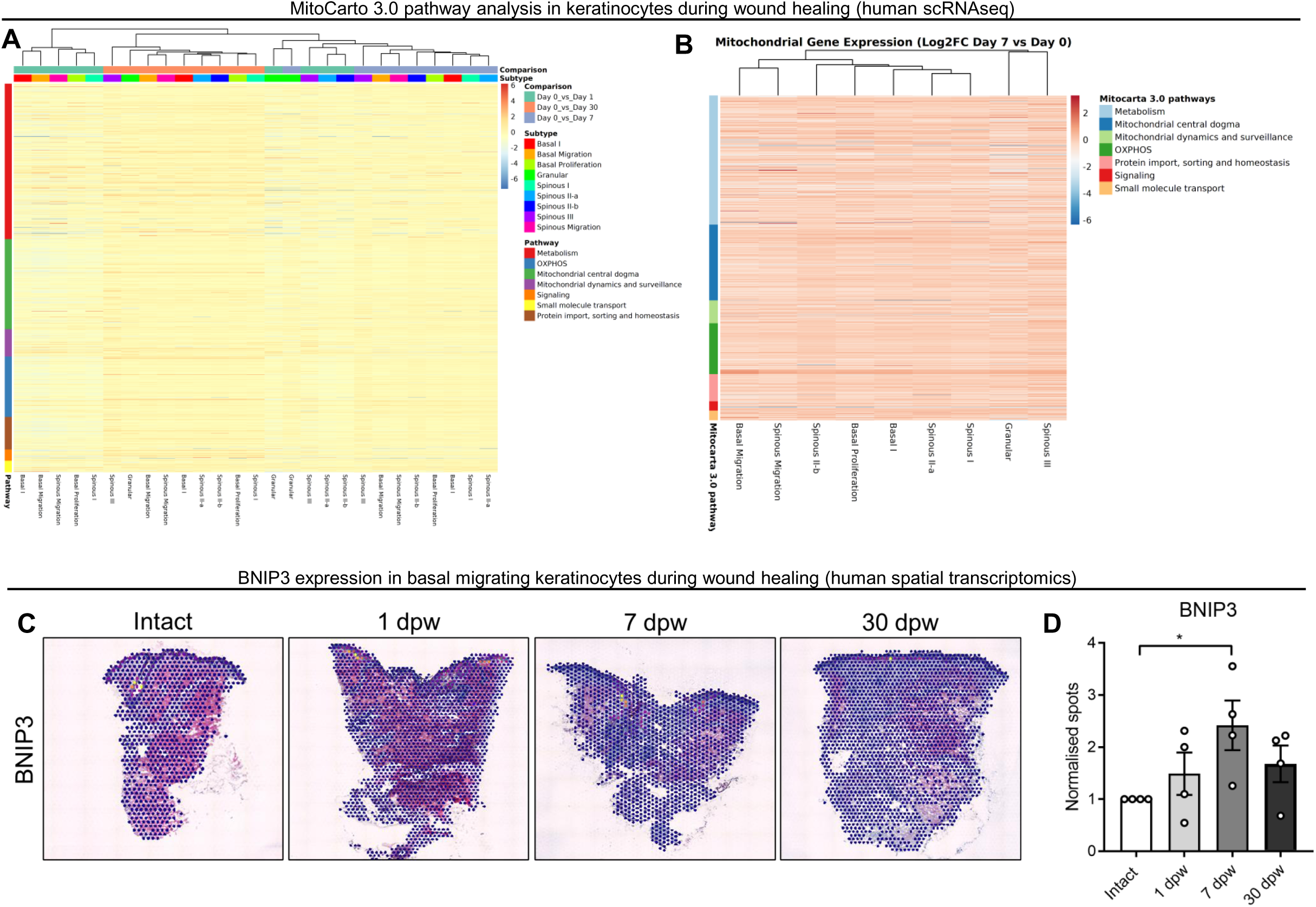
Mitochondrial pathway and *BNIP3* analysis in human wound healing. (**A-B**) Heat map depicting the log2fold change in mitochondria related genes in (**A**) all keratinocyte subtypes at days 1, 7, and 30 vs day 0, and (**B**) at day 7 vs day 0. (**C-D**) Representtive images and (**D**) quantification (mean ± SEM) of BNIP3 spots normalised to intact from spatial transcriptomics data of human wound healing.

**Supplementary Figure 2.**
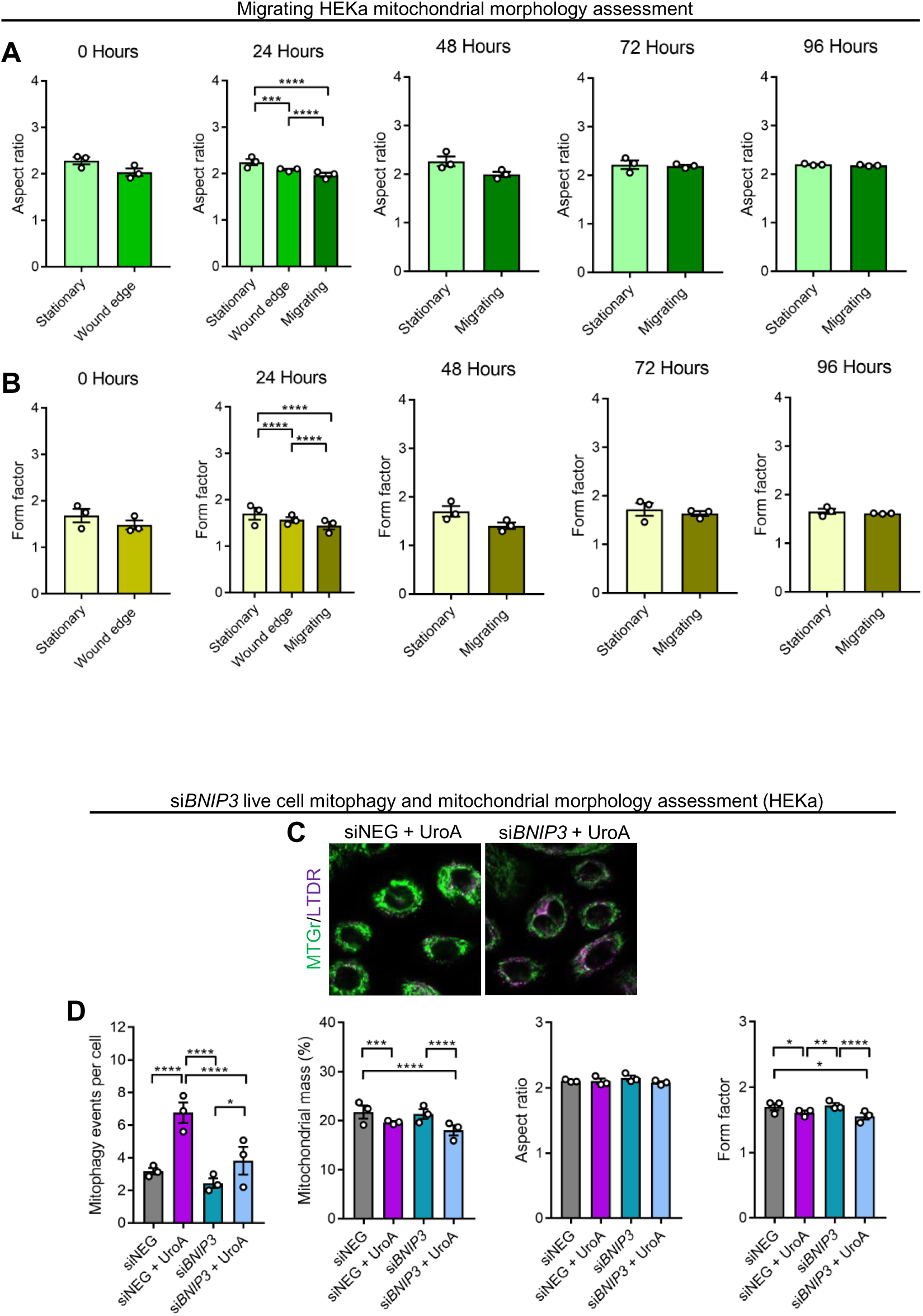
Live cell mitophagy and mitochondrial morphology analysis in *BNIP3* KD HEKA. (**A-B**) Quantification (mean ± SEM) of mitochondrial aspect ratio and form factor in WT HEKa. Two-way ANOVA or students’s t-test. **** = p < 0.0001; *** = p < 0.001. N = 30 cells in 3 separate biological replicates within each condition. (**C**) Representative fluorescence images of MTGr and LTDR staining in WT siNEG and si*BNIP3* HEKa. (**D**) Quantification (mean ± SEM) of mitophagy events per cell, mitochondrial mass, aspect ratio, and form factor in WT HEKa. Two-way ANOVA, **** = p < 0.0001; *** = p < 0.001; ** = p < 0.01, * = p < 0.05. N = 15 cells in 3 separate biological replicates within each condition.

**Supplementary Figure 3.**
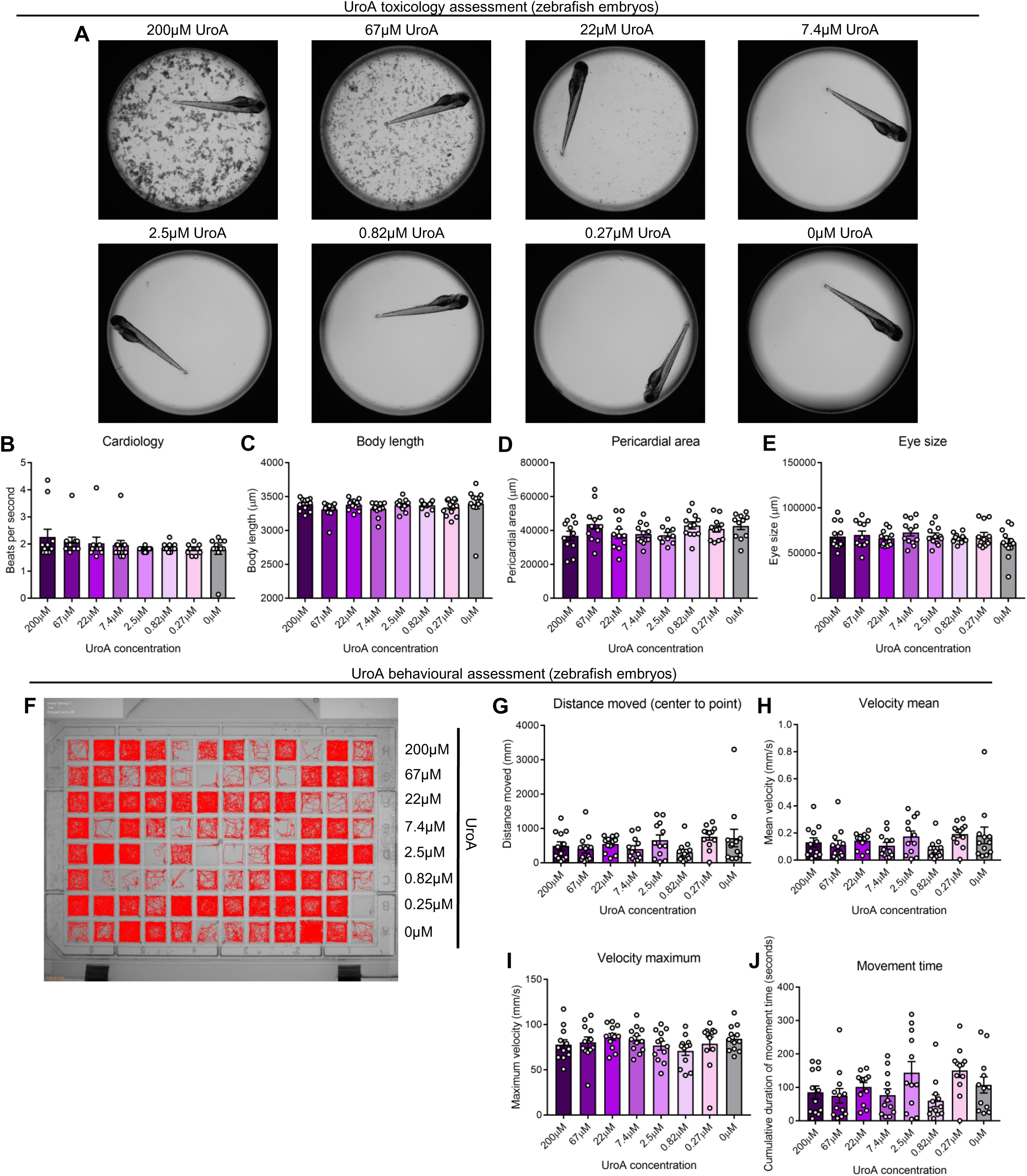
Live cell mitophagy and mitochondrial morphology analysis in *BNIP3* KD HEKA. (**A**) Representative brightfield images of zebrafish embryos with treatment with various concentration of UroA. (**B-E**) Quantification (mean ± SEM) of (**B**) cardiology, (**C**) body length (µm), (**C**) pericardial area (µm), and (**E**) eye size (µm). (**F**) Rrepresentative tracking map of zebrafish embryos over 1 hour. (**G-J**) Quantification (mean ± SEM) of (**G**) distance moved from centre to periphery, (**H**) mean velocity (mm/s), (**I**) maximum velocity (mm/s), and (**J**) movement time (seconds).

**Supplementary Figure 4.**
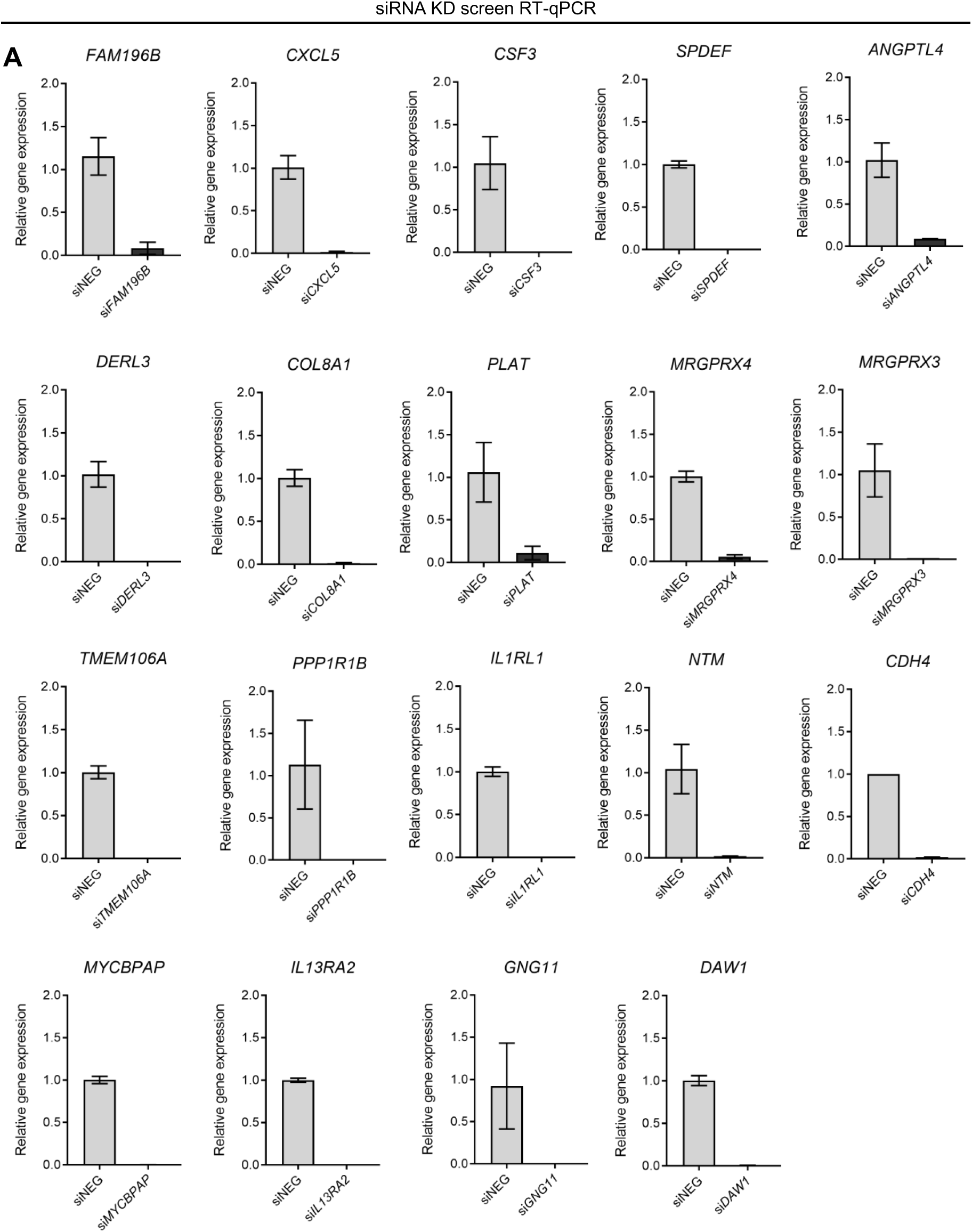
Validation of gene expression in siRNA screen. (**A**) Quantification (mean ± SEM) of gene expression of the top 20 significantly upregulated genes (from RNAseq) following siRNA KD.

**Supplementary Figure 5.**
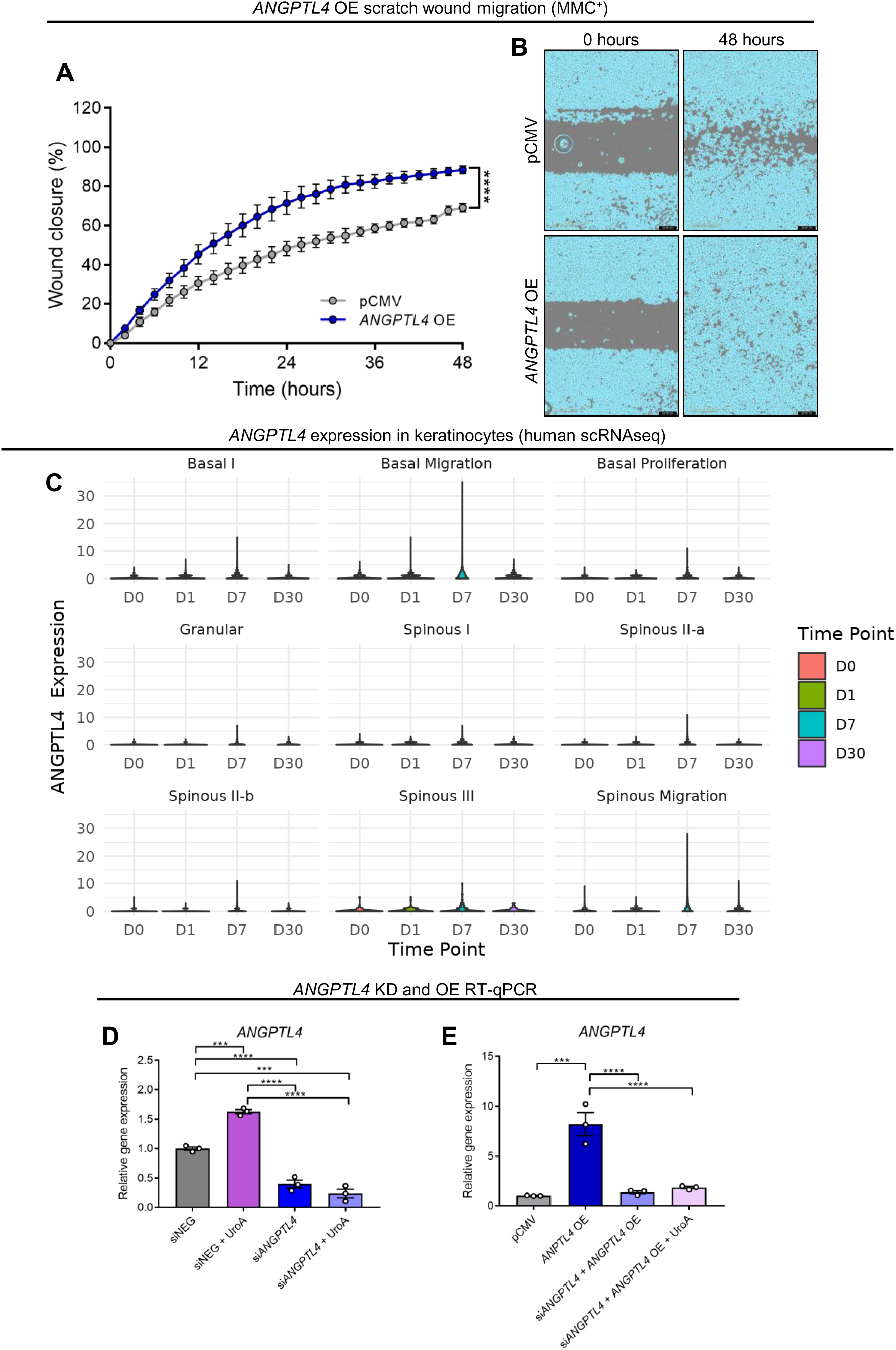
ANGPTL4 is upregulated at the proliferative stage of wound healing and accelerates keratinocyte migration *in vitro*. (**A-B**) Quantifiacation (mean ± SEM) and (**B**) representative images of scratch wound migration assessment in *ANGPTLL4* OE HEKa. Students t-test **** = p < 0.0001. Each dot represents the mean of an individual biological replicate (n =3) containing at least 3 technical repeats. (**C**) Violin plot depicting *ANGPTL4* epression in keratinocyte subtypes from scRNAseq analysis of human wound healing. (**D-E**) Quantification (mean ± SEM) of ANGPTL4 gene epression. Two-way ANOVA **** = p < 0.0001

**Supplementary Figure 6.**
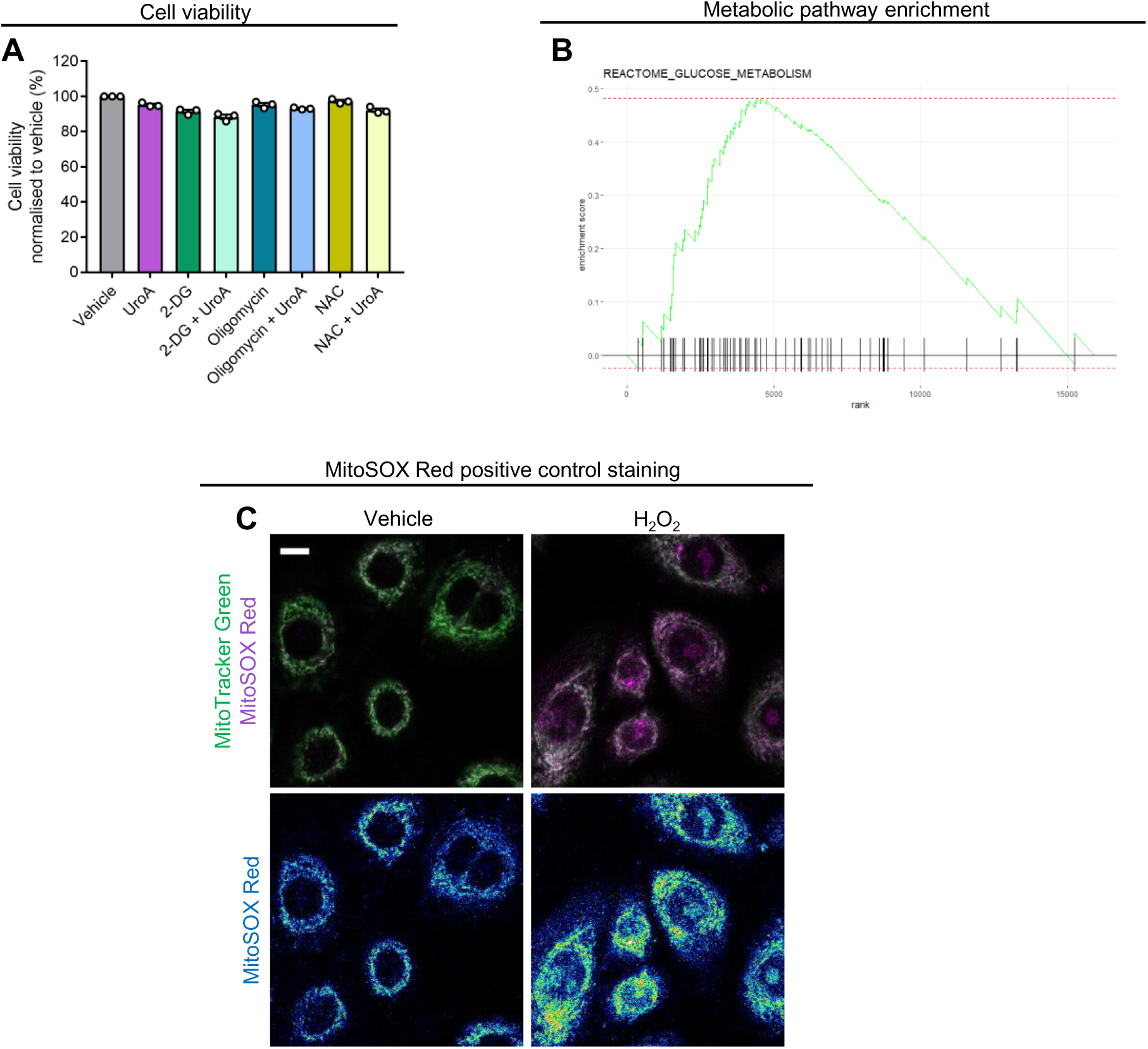
UroA upregulates glycolytic metabolism in keratinocytes. (**A**) Quantification (mean ± SEM) of cell viability. Two-way ANOVA. Each dot represents the mean of an individual biological replicate (n = 3) containing at least 3 technical repeats. (**B**) Enrichment plot for glucose metabolism. (**C**) Representative fluorescence images and (**H**) quantification (mean ± SEM) of MitoTracker Green and MitoSOX Red live cell staining in HEKa.

**Supplementary Figure 7.**
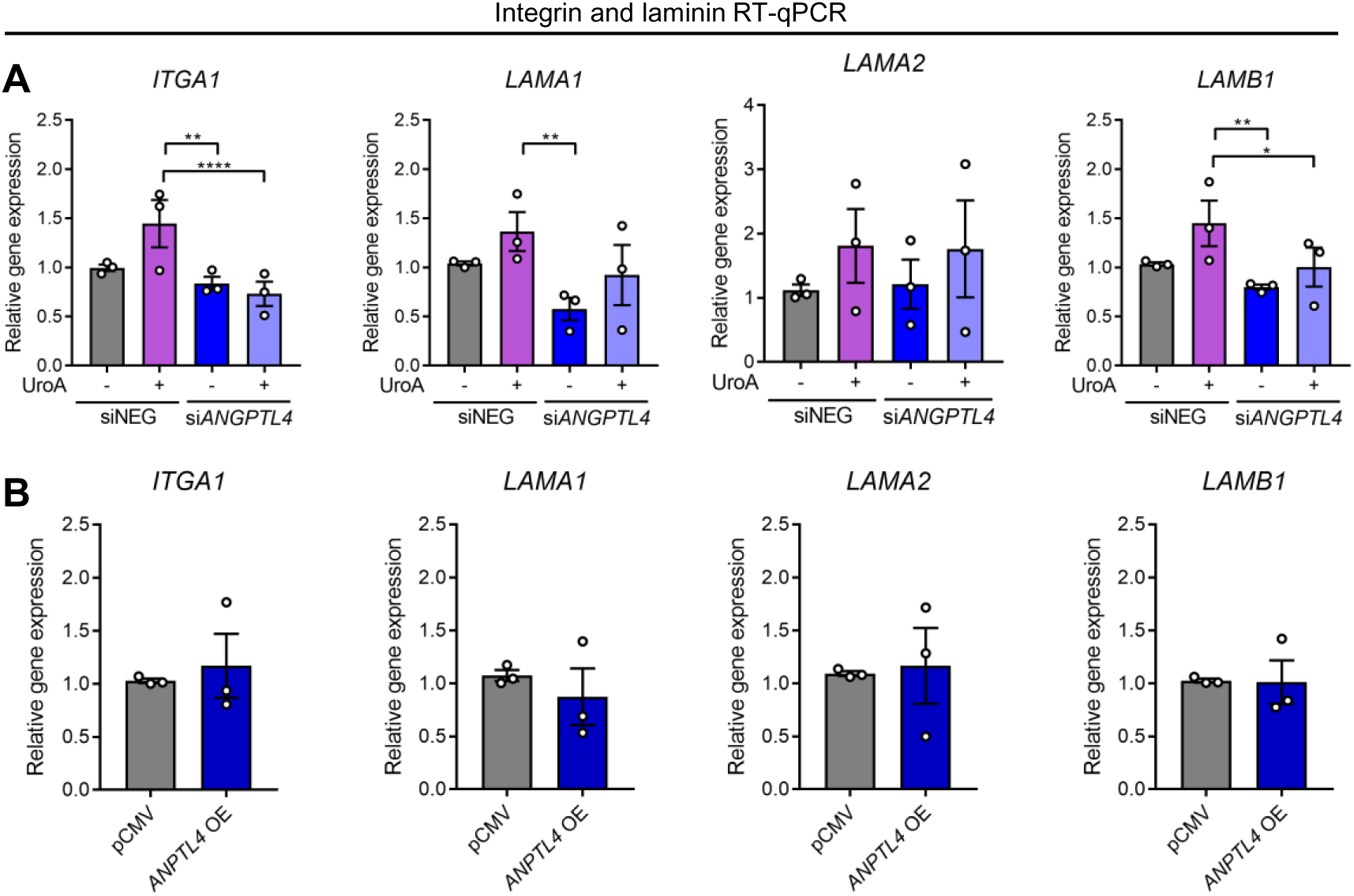
Assessment of laminin and integrin gene expression. (**A-B**) Quantification (mean ± SEM) of integrin and laminin gene expression in (**A**) si*ANGPT4* and (**B**) *ANGPTL4* OE HEKa. Two-way ANOVA and students t-test. **** = p < 0.0001, ** = p < 0.01, * = p < 0.05.

