## Supplemental Figures for "Mitophagy promotes metabolic reprogramming to enhance keratinocyte migration via ANGPTL4 during wound healing"

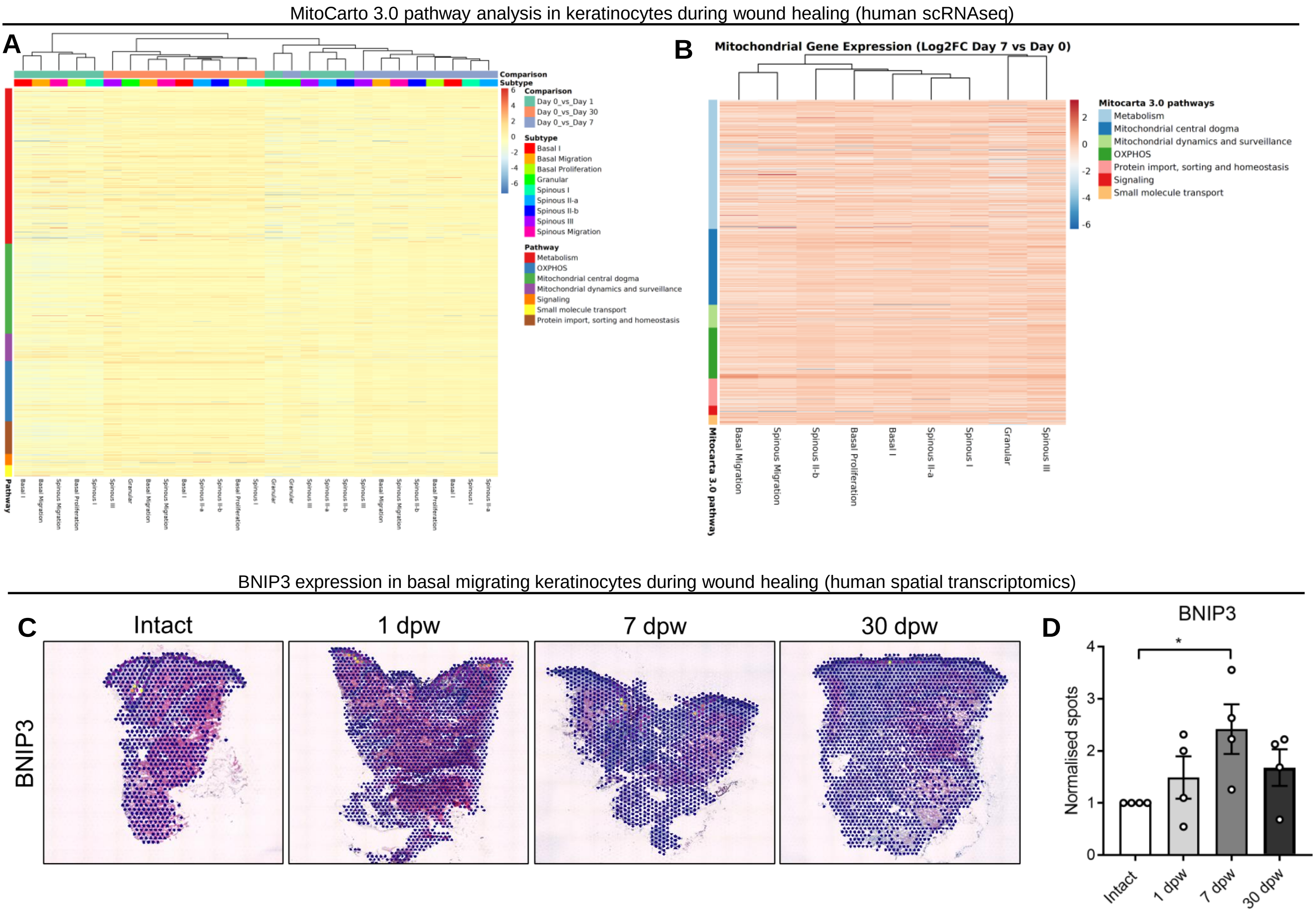


**Supplementary Figure 1 – Mitochondrial pathway and *BNIP3* analysis in human wound healing.**
(**A-B**) Heat map depicting the log2fold change in mitochondria related genes in (**A**) all keratinocyte subtypes at days 1, 7, and 30 vs day 0, and (**B**) at day 7 vs day 0. (**C-D**) Representative images and (**D**) quantification (mean ± SEM) of BNIP3 spots normalised to intact from spatial transcriptomics data of human wound healing.

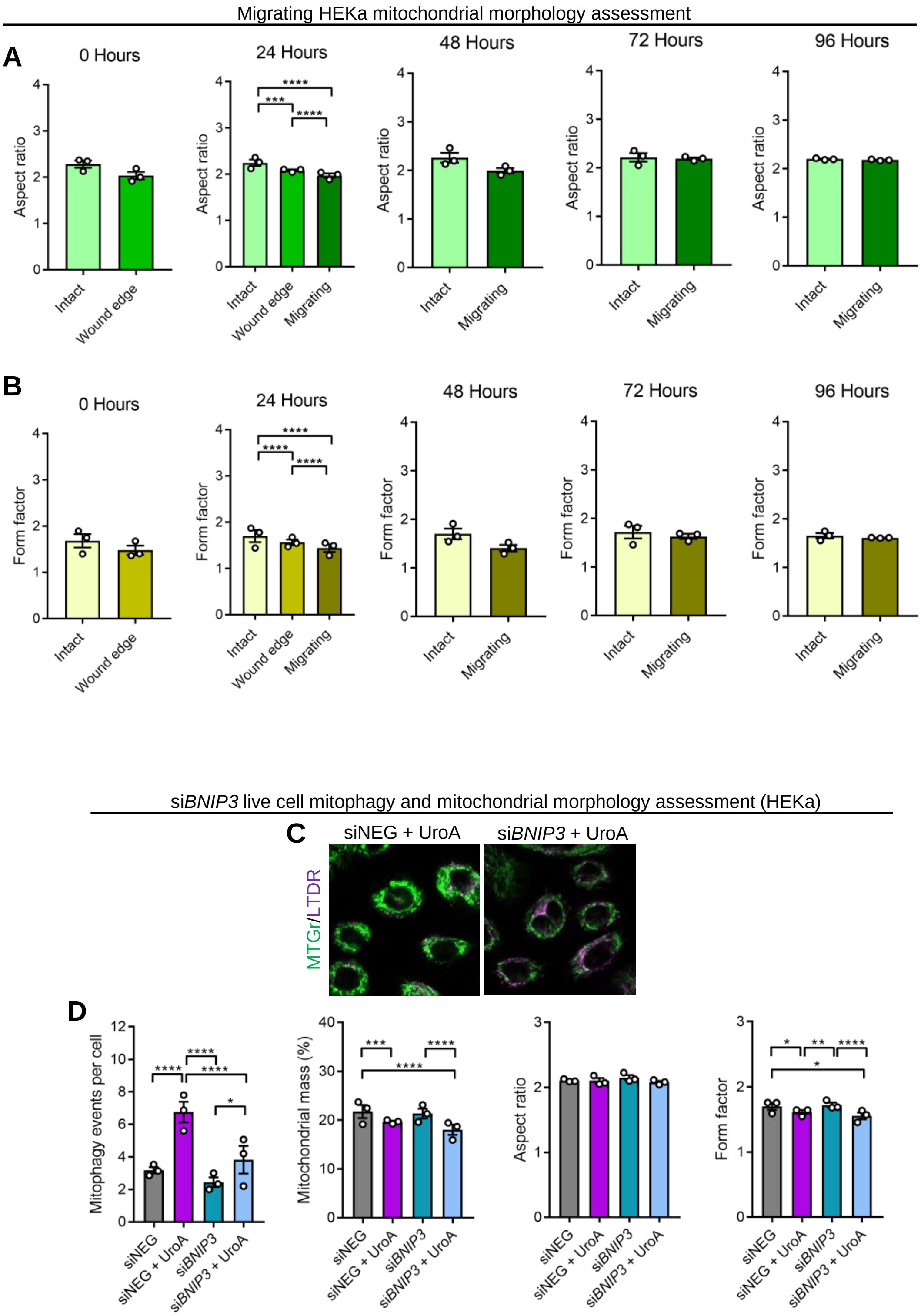


**Supplementary Figure 2 – Live cell mitophagy and mitochondrial morphology analysis in *BNIP3* KD HEKA.** (**A-B**) Quantification (mean ± SEM) of mitochondrial aspect ratio and form factor in WT HEKa. Two-way ANOVA or student's t-test. **** = $p < 0.0001$; *** = $p < 0.001$. N = 30 cells in 3 separate biological replicates within each condition. (**C**) Representative fluorescence images of MTGr and LTDR staining in WT siNEG and si*BNIP3* HEKa. (**D**) Quantification (mean ± SEM) of mitophagy events per cell, mitochondrial mass, aspect ratio, and form factor in WT HEKa. Two-way ANOVA, **** = $p < 0.0001$; *** = $p < 0.001$; ** = $p < 0.01$, * = $p < 0.05$. N = 15 cells in 3 separate biological replicates within each condition.

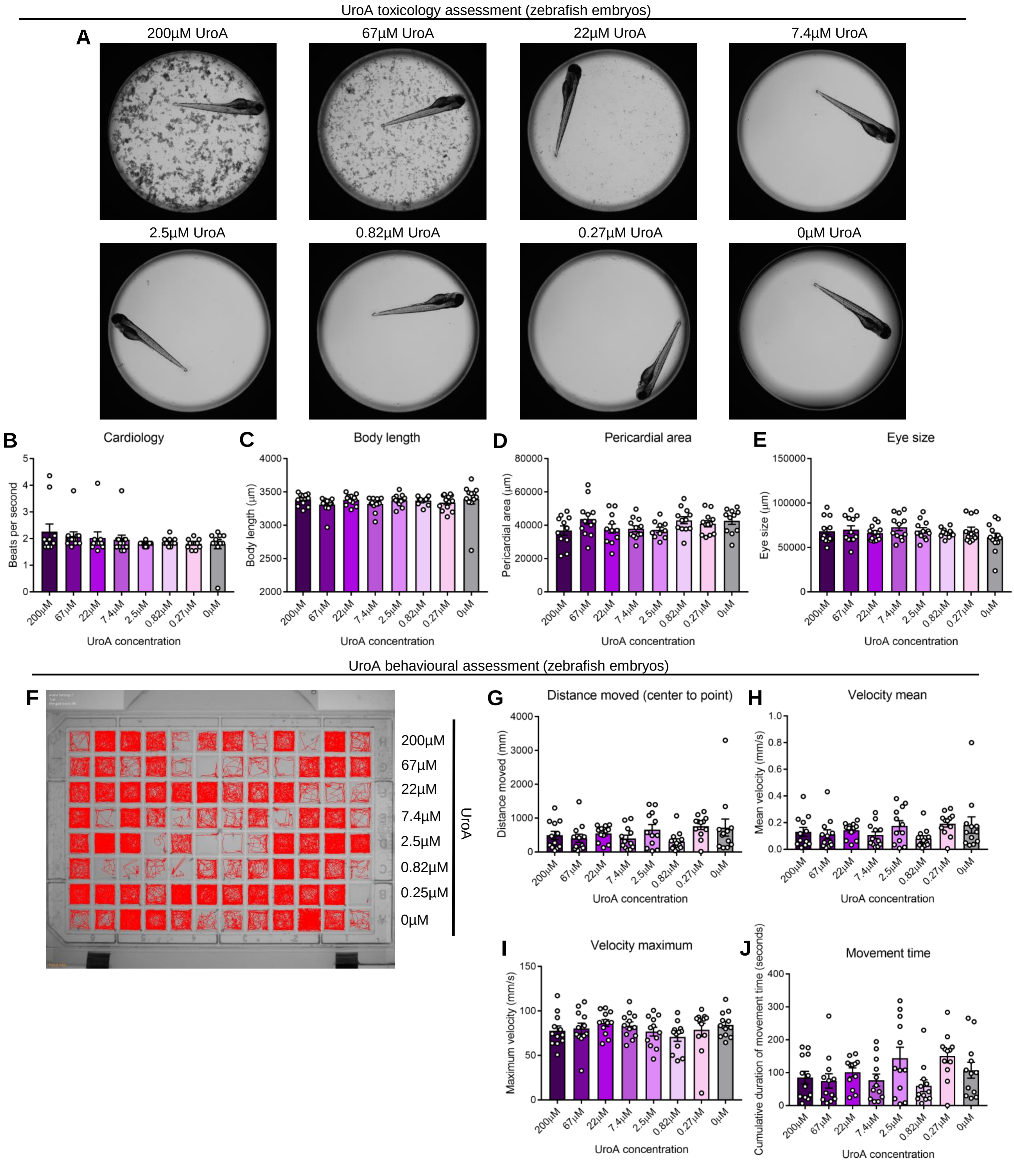


**Supplementary Figure 3 – Analysis of UroA toxicity in zebrafish embryo model.**
(**A**) Representative brightfield images of zebrafish embryos with treatment with various concentration of UroA. (**B-E**) Quantification (mean ± SEM) of (**B**) cardiology, (**C**) body length (µm), (**C**) pericardial area (µm), and (**E**) eye size (µm). (**F**) A tracking map of zebrafish embryos over 1 hour. (**G-J**)
Quantification (mean ± SEM) of (**G**) distance moved from centre to periphery, (**H**) mean velocity (mm/s), (**I**) maximum velocity (mm/s), and (**J**) movement time (seconds).

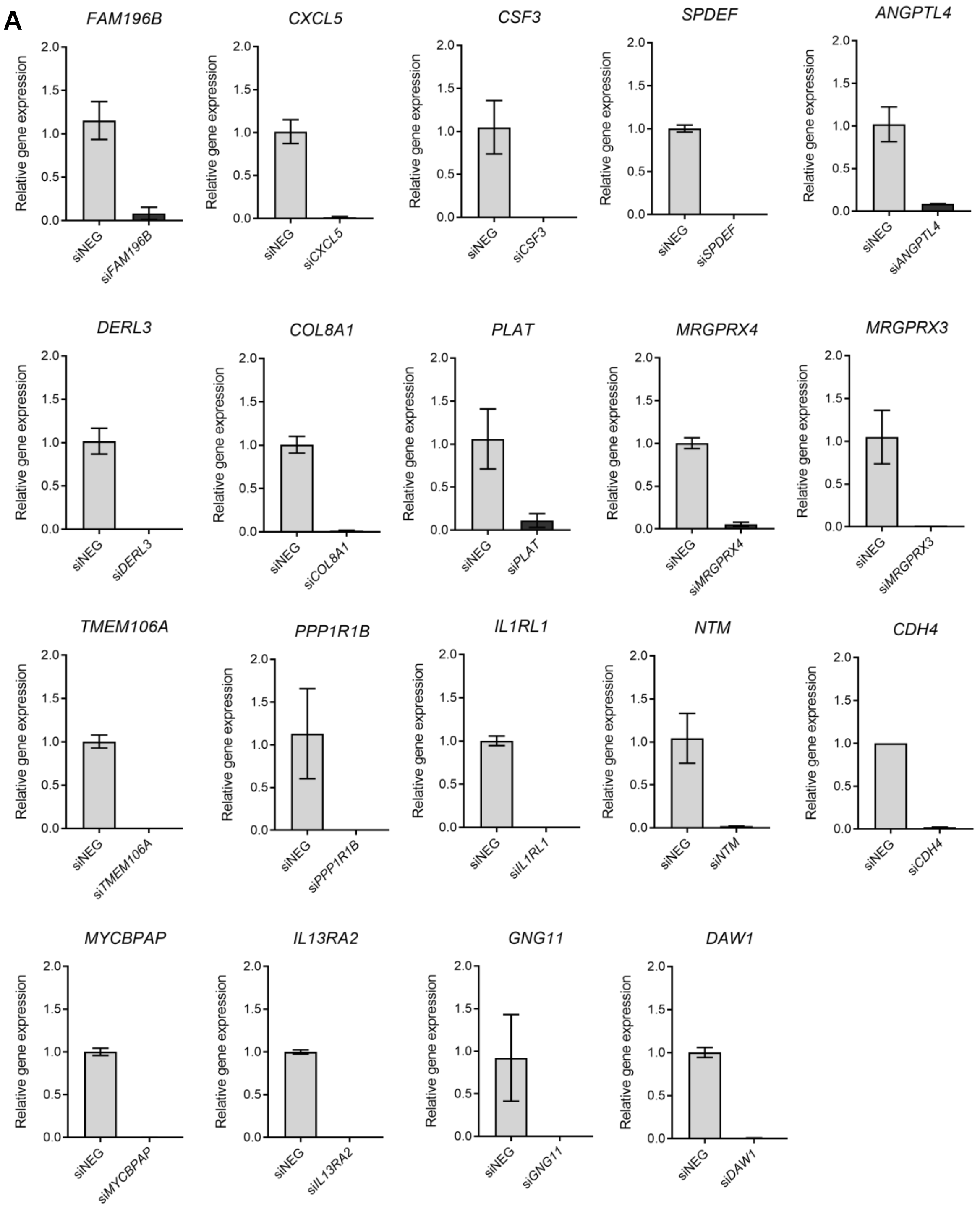


**Supplementary Figure 4 – Validation of gene expression in siRNA screen.**
(**A**) Quantification (mean ± SEM) of gene expression of the top 20 significantly upregulated genes (from RNAseq) following siRNA KD.

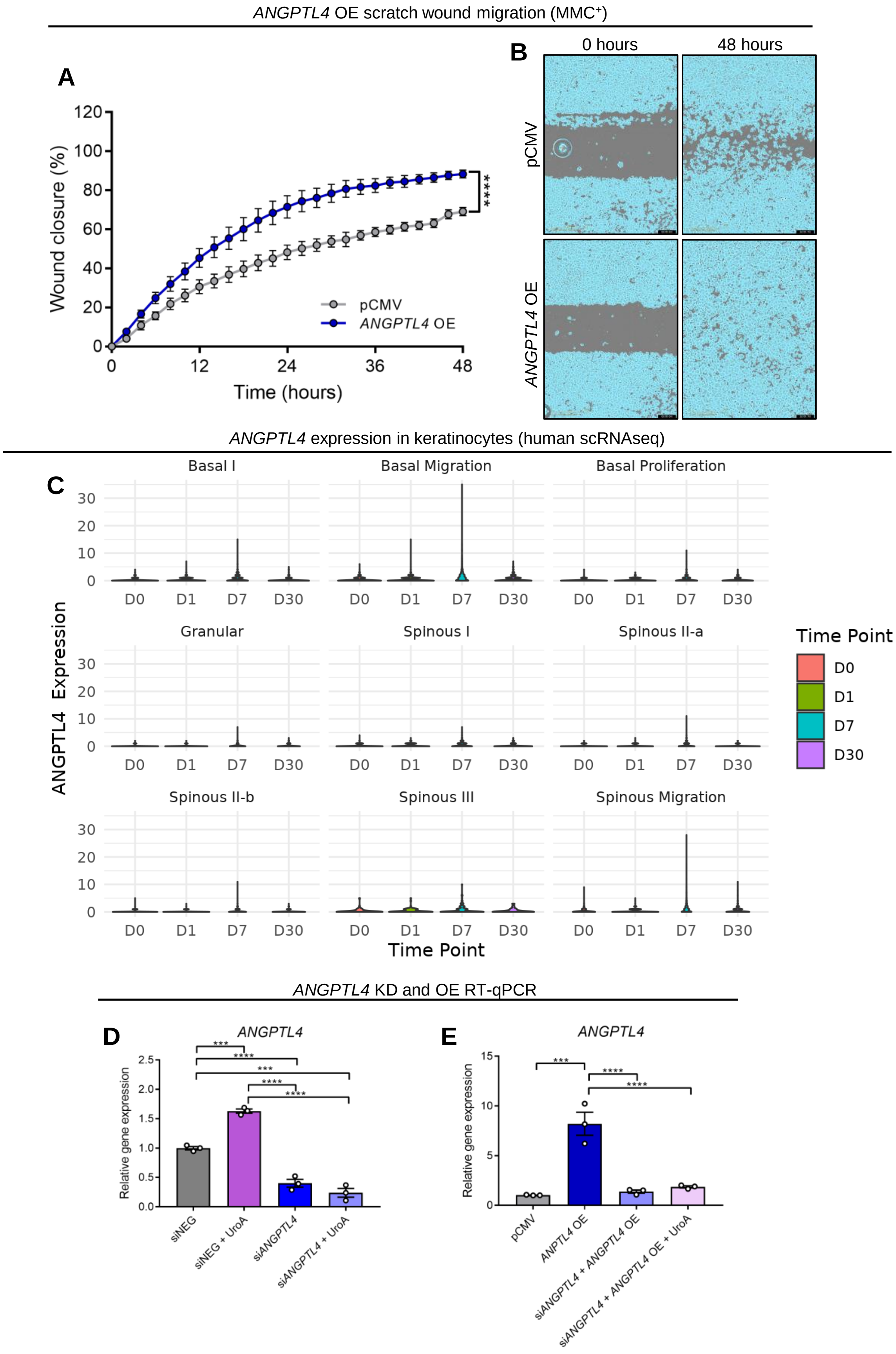


**Supplementary Figure 5 – ANGPTL4 is upregulated at the proliferative stage of wound healing and accelerates keratinocyte migration *in vitro.***

(**A-B**) Quantifiacation (mean ± SEM) and (**B**) representative images of scratch wound migration assessment in *ANGPTLL4* OE HEKa. Students t-test **** = $p < 0.0001$. Each dot represents the mean of an individual biological replicate (n =3) containing at least 3 technical repeats. (**C**) Violin plot depicting *ANGPTL4* epression in keratinocyte subtypes from scRNAseq analysis of human wound healing. (**D-E**) Quantification (mean ± SEM) of ANGPTL4 gene epression. Two-way ANOVA **** = $p < 0.0001$

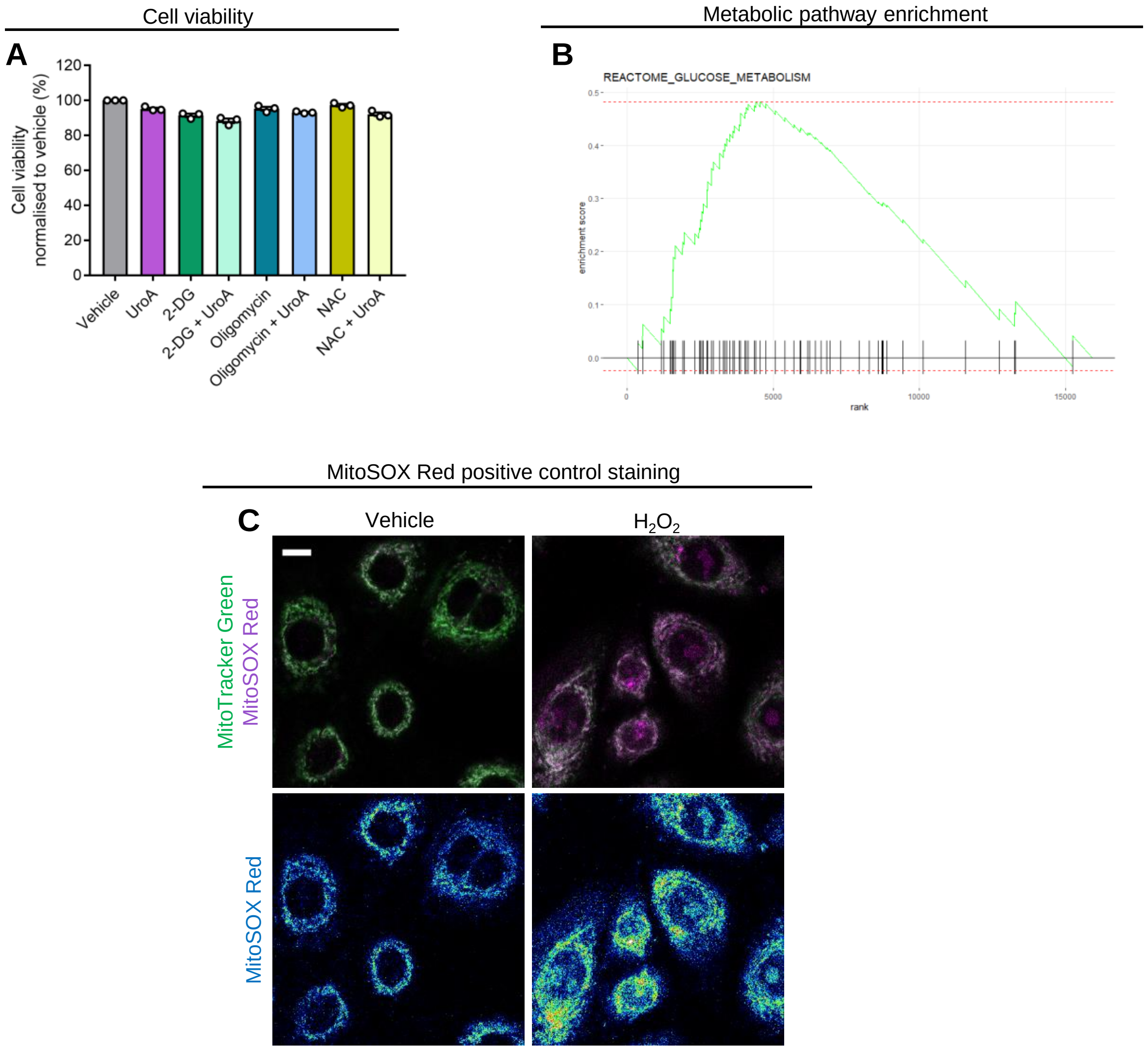


**Supplementary Figure 6 – UroA upregulates glycolytic metabolism in keratinocytes.**
(**A**) Quantification (mean ± SEM) of cell viability. Two-way ANOVA. Each dot represents the mean of an individual biological replicate (n = 3) containing at least 3 technical repeats. (**B**) Enrichment plot for glucose metabolism. (**C**) Representative fluorescence images and (**H**) quantification (mean ± SEM) of MitoTracker Green and MitoSOX Red live cell staining in HEKa.

Integrin and laminin RT-qPCR

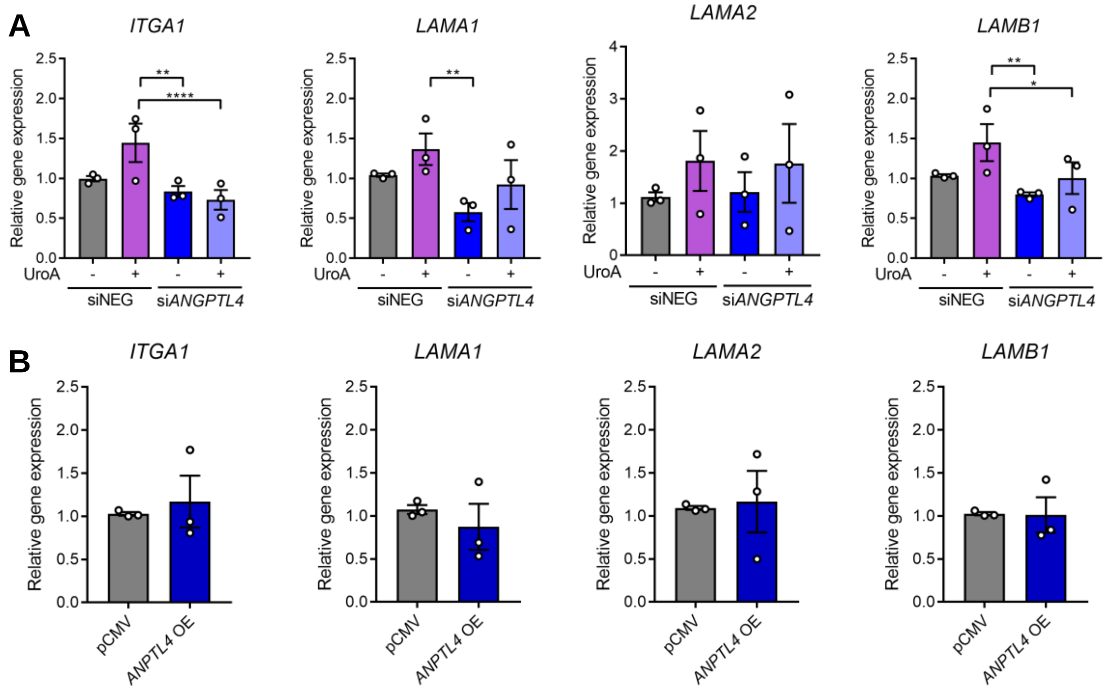


**Supplementary Figure 7 – Assessment of laminin and integrin gene expression.**
(**A-B**) Quantification (mean ± SEM) of integrin and laminin gene expression in (**A**) si*ANGPT4* and (**B**) *ANGPTL4* OE HEKa. Two-way ANOVA and students t-test. **** = $p < 0.0001$, ** = $p < 0.01$, * = $p < 0.05$**.**
